# A durably colonizing engineered native symbiont enables sustained intestinal delivery for metabolic dysfunction and colitis

**DOI:** 10.64898/2026.08.02.741375

**Authors:** Qixiang Zhao, Yong Ding, Sen Yan, Haohan Ma, Yicun Wang, Siqi Guo, Xi Luo, Yanli Pang, Changtao Jiang, Kai Wang

**Affiliations:** Department of Physiology and Pathophysiology, Center for Obesity and Metabolic Disease Research, State Key Laboratory of Vascular Homeostasis and Remodeling, School of Basic Medical Sciences, Peking University, Beijing 100191, China; Department of Immunology, School of Basic Medical Sciences, NHC Key Laboratory of Medical Immunology, Peking University, Beijing 100191, China; Institute of Advanced Clinical Medicine, State Key Laboratory of Female Fertility Promotion, Center for Reproductive Medicine, Third Hospital, Peking University, Beijing, China; National Clinical Research Center for Obstetrics and Gynecology (Peking University Third Hospital), Beijing, China; Beijing Key Laboratory of Reproductive Endocrinology and Assisted Reproductive Technology, Beijing, China; Center of Basic Medical Research, Institute of Medical Innovation and Research, Peking University Third Hospital, Beijing, China

**Keywords:** Native probiotics, engineering bacteria, persistent colonization, microbial therapy, nicotinic acid, metabolic disorder, IL-10, colitis

## Abstract

**Background:** Engineered bacterial therapeutics represent a promising strategy for sustained intestinal delivery of therapeutic molecules, but their efficacy is limited by inefficient colonisation, safety concerns and the need for repeated administration or auxiliary delivery systems.

**Objective:** To develop a safety-optimised native bacterial chassis capable of long-term gut colonisation and sustained therapeutic delivery for intestinal inflammatory and metabolic diseases.

**Design:** Native murine *Escherichia coli* isolates were screened for antibiotic susceptibility, genetic tractability and long-term intestinal colonisation. The selected strain, MEc30, was further optimised by deleting the putative virulence-associated *clb* and *irp* loci. MEc30 was then engineered to deliver murine interleukin-10 (MEc30-mIL-10) or produce nicotinic acid (MEc30-NA), and therapeutic efficacy was evaluated in *Il10*^-/-^ colitis and high-fat diet-induced metabolic dysfunction models.

**Results:** MEc30 achieved stable lifelong colonisation of the murine intestine after a single oral administration, without antibiotic preconditioning or auxiliary delivery systems, and did not detectably disturb host physiology or gut microbial ecology. Deletion of *clb* and *irp* abolished potential colibactin- and yersiniabactin-associated biosafety risks while preserving bacterial growth and colonisation capacity. MEc30-NA continuously produced nicotinic acid in the gut, activated epithelial GPR109a-associated barrier signalling, improved glucose and lipid metabolism, reduced systemic inflammation and avoided the sharp peak exposure associated with conventional nicotinic acid administration. MEc30-mIL-10 enabled sustained intestinal IL-10 delivery, suppressed inflammatory macrophage activation, improved barrier integrity and alleviated spontaneous colitis in *Il10*^-/-^ mice.

**Conclusion:** This study identifies MEc30 as a durable and safety-optimised native *E. coli* chassis for sustained intestinal therapeutic delivery. Engineered native symbionts may provide a long-acting live biotherapeutic strategy for chronic intestinal inflammatory and metabolic diseases.

## INTRODUCTION

Intestinal microorganisms have co-evolved with their hosts and are central regulators of metabolism, mucosal immunity and tissue homeostasis. Microbial metabolites, including short-chain fatty acids, vitamins, amino acid-derived products and secondary bile acids, regulate epithelial barrier integrity, immune responses and systemic physiology^1–3^. These host-microbe interactions provide a biological rationale for using engineered bacteria as living therapeutics capable of producing defined molecules directly within the gut. However, the therapeutic performance of current engineered microbes is often limited by inefficient intestinal colonisation, variable persistence across hosts, the need for repeated administration or antibiotic preconditioning, and potential biosafety concerns^4^ ^5^.

Native gut microorganisms may provide a solution to these limitations because they are adapted to the intestinal ecosystem and can establish stable host-associated relationships. Previous studies have shown that host-derived native *Escherichia coli* strains, including MP-1, NGF-1 and EcAZ-2, can achieve prolonged intestinal colonisation and mediate sustained physiological effects after engineering^6–8^. Nevertheless, the therapeutic potential of native intestinal isolates as chassis for disease intervention remains insufficiently explored. Systematic strategies for their selection, safety evaluation, and optimization are still poorly defined. Moreover, native gut bacteria may carry antibiotic resistance genes or putative virulence-associated loci, raising concerns regarding their long-term safety in therapeutic settings^9^.

Here, we isolated and screened native murine *E. coli* strains for antibiotic susceptibility, genetic tractability and long-term intestinal colonisation. We identified MEc30 as a rare native strain capable of stable, lifelong colonisation of the murine gut after a single oral administration without antibiotic preconditioning or auxiliary delivery systems. We further optimised this chassis by deleting the putative virulence-associated *clb* and *irp* loci, which encode colibactin- and yersiniabactin-associated functions, respectively^10^. Using this safety-optimised native chassis, we generated MEc30-NA for continuous nicotinic acid production and MEc30-mIL-10 for sustained intestinal IL-10 delivery. These engineered strains alleviated high-fat diet (HFD)-induced metabolic dysfunction and IL-10-deficiency-associated colitis, respectively, supporting engineered native symbionts as a durable live biotherapeutic strategy for chronic intestinal inflammatory and metabolic diseases.

## RESULTS

### Isolation, genetic manipulation, and colonisation screening of native *E. coli*

*E*. *coli* Nissle 1917 (EcN) is a widely utilized bacterial chassis known for its versatile metabolic adaptability in engineered probiotic therapies^11^. However, the efficacy of colonisation by EcN varies significantly across different hosts and environmental conditions, with some studies reporting that stable colonisation requires pre-treatment with antibiotics to clear native intestinal microbiota, limiting its practical therapeutic utility^12^ ^13^.

Russell and colleagues previously explored a host-derived native bacterial chassis strategy, demonstrating that intestinal transgene delivery with native *E. coli* chassis can enable persistent physiological changes *in vivo*^6^. To develop alternative native bacterial chassis with superior colonisation properties for the treatment of genetic and metabolic disorders, we isolated 10 native *E*. *coli* strains from 5 murine guts. Genetic manipulation of undomesticated strains is often challenging due to factors such as restriction-modification (RM) systems, making genome editability an important consideration when selecting an engineering bacterial chassis^14^. Accordingly, we employed a dual-plasmid system, pEcCas/pEcgRNA^15^, to evaluate the genome editability of isolated native *E*. *coli* strains. Due to intrinsic antibiotic resistance, GFP insertion at the *exo* locus was successfully achieved in only two strains. *In vivo* colonisation analysis showed that none of these initially isolated native strains maintained detectable colonisation beyond 30 days, indicating that durable colonisation is not a general property of native *E. coli* isolates (Figures 1A-C).

**Figure 1.**
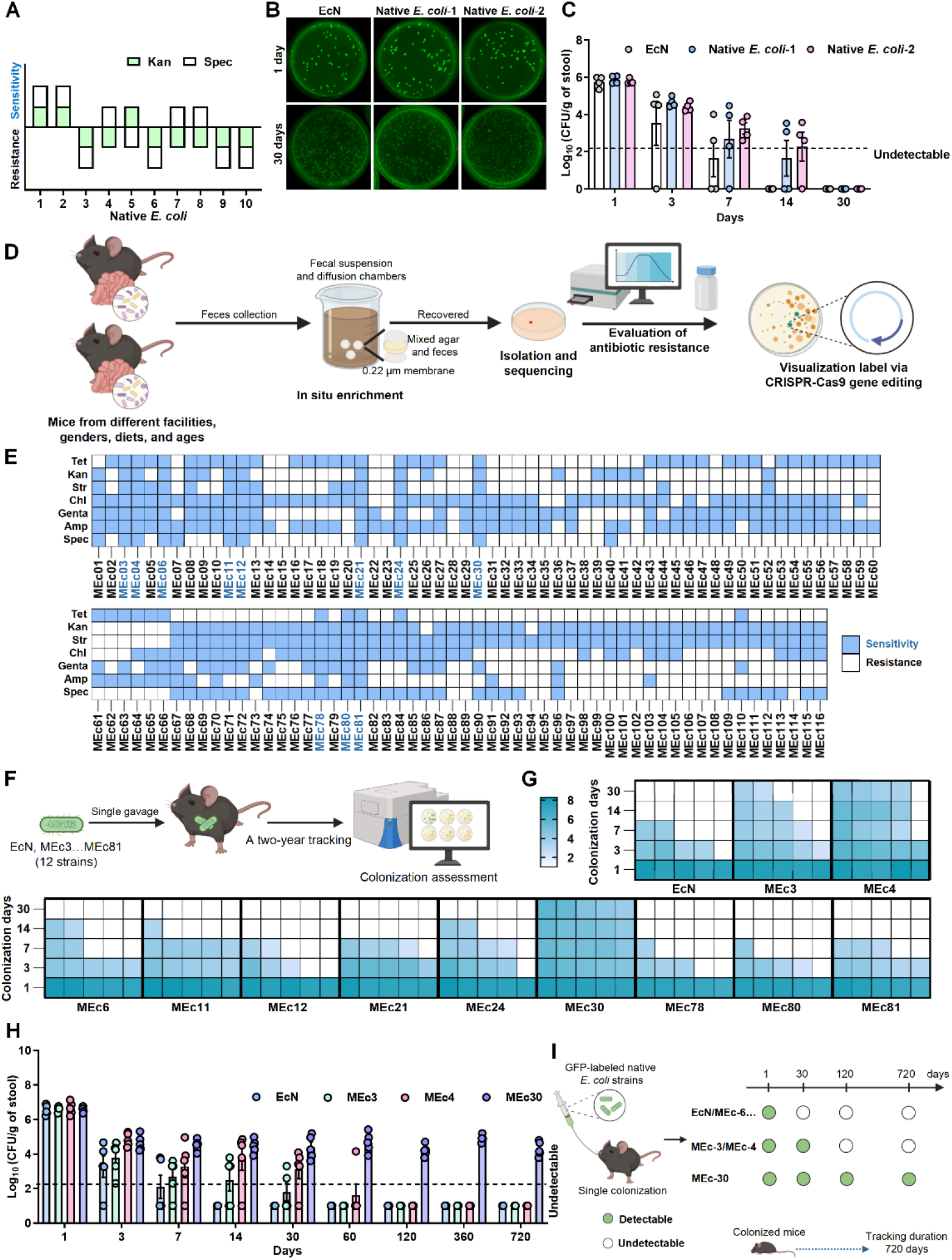
Isolation and colonisation screening of native *E. coli* from the murine gut. **(A)** Antibiotic resistance and sensitivity profiles of 10 native *E. coli* isolates against kanamycin (Kan) and spectinomycin (Spec). Bars above the axis indicate sensitivity, while bars below indicate resistance. **(B)** Representative images of GFP-positive colonies from fecal samples of *Escherichia coli* Nissle 1917 (EcN), Native *E. coli*-1, and Native *E. coli*-2 at 1 day and 30 days. All plates were imaged under 488 nm excitation. **(C)** Quantification of fecal colony forming unit (CFU) counts for persistent strains (EcN, native *E*. *coli*-1, or native *E*. *coli*-2) at 1 day,3 days, 7 days, 14 days, and 30 days post single administration. **(D)** Schematic overview of the workflow for isolating and characterizing native intestinal *E. coli*. Fecal samples collected from mice of different facilities, genders, diets, and ages were subjected to *in situ* enrichment using diffusion chambers to promote the recovery of native gut bacteria. Enriched samples were plated for *E. coli* isolation and verified by 16S rRNA gene sequencing. Each isolate was systematically evaluated for antibiotic susceptibility against seven commonly used antibiotics. Candidate strains with broad antibiotic sensitivity were subsequently engineered to stably express GFP via CRISPR-Cas9-mediated chromosomal integration, enabling in vivo visualization and colonisation tracking. **(E)** Antibiotic resistance profiles of 116 native *E. coli* isolates against seven antibiotics (Spectinomycin, Ampicillin, Gentamicin, Chloramphenicol, Streptomycin, Kanamycin, Tetracycline). Strains highlighted in blue (MEc3, MEc4, MEc6, MEc11, MEc12, MEc21, MEc24, MEc30, MEc78, MEc80, MEc81) were broadly sensitive and selected for subsequent genome editing. **(F)** Schematic overview of the long-term colonisation assessment of native *E. coli* strains. Twelve representative isolates (EcN, MEc3, MEc4, MEc6, MEc11, MEc12, MEc21, MEc24, MEc30, MEc78, MEc80, MEc81) were administered to mice via a single oral gavage. Colonisation dynamics were monitored over two year. Fecal bacterial loads were quantified by plating and expressed as log_10_ (CFU/g of stool). **(G)** Heatmap showing fecal CFU counts for EcN and 11 native *E. coli* strains at multiple timepoints within 30 days post single gavage (1, 3, 7, 14, and 30 days). **(H)** Quantification of fecal CFU counts for persistent strains (EcN, MEc3, MEc4, MEc30) over an extended time course, including days 1, 3, 7, 14, 30, 60, 120, 360, and 720 (approximating the average lifespan of laboratory mice). **(I)** Schematic summary of the long-term colonisation outcomes of representative GFP-labeled native *E. coli* strains after a single oral administration. Strains such as EcN, MEc6, and other non-persistent isolates were detectable only at early time points, MEc3 and MEc4 displayed intermediate persistence up to 30 days, whereas MEc30 remained detectable through 720 days, indicating durable long-term colonization. All data are presented as mean ± SEM. *n* = 4 in each group in (C), *n* = 5 in each group in (G)-(H).

Building on our previous finding that *in situ* cultivation can enrich intestinal microbes^16^ and may better preserve the ecological competitiveness of gut-resident isolates^17^, we explored this strategy to mimic gut environment for native microbes’ enrichment. Fecal samples were collected from mice across different housing rooms, sexes, diets and ages, followed by *in situ* fecal cultivation and recovery of native *E. coli* isolates. In total, 116 native *E. coli* isolates were obtained and evaluated for antibiotic susceptibility and genetic tractability (Figures 1D and 1E). Approximately 90% of these isolates showed resistance to common antibiotics, whereas only 11 strains were broadly antibiotic-sensitive and therefore selected as candidate engineering chassis. These 11 strains were successfully labelled by chromosomal GFP insertion, and GFP expression did not affect bacterial growth (Figures S1A-C).

Because intestinal persistence is a key determinant of efficacy for live bacterial therapeutics, we next assessed the colonisation capacity of these candidate chassis strains after a single oral administration to 8-week-old C57BL/6J mice without antibiotic pretreatment (Figure 1F). All strains were detectable in feces at day 1, but their persistence rapidly diverged over time. By day 30, most strains, including EcN, had markedly declined or become undetectable, whereas MEc3, MEc4 and MEc30 remained detectable (Figure 1G and Figure S2A). qPCR-based quantification confirmed the GFP-based tracking results (Figures S2B-D). After 120 days, MEc30 was the only strain that maintained stable intestinal colonisation (Figure 1H). Remarkably, MEc30 remained detectable up to 720 days after a single gavage, a duration approximating the lifespan of C57BL/6J mice (Figures 1H and 1I; Figures S2E and S2F). These findings identify MEc30 as a rare genetically tractable native *E. coli* strain with durable, lifelong intestinal colonisation capacity, supporting its use as a chassis for long-term therapeutic engineering.

### Systematic profiling of MEc30 colonisation stability and host safety

To further define the colonisation robustness and biosafety of MEc30 as a candidate native bacterial chassis, we next examined whether its engraftment was influenced by inoculation dose or host physiological conditions, and assessed its spatial distribution, host compatibility, and biosafety *in vivo*. Host-native *E. coli* failed to establish durable engraftment when the inoculum was below 1×10^6^ CFU, we found that the colonisation efficiency of MEc30 declined significantly when the dose was below this threshold (Figure S3A-C). Because gut microbiota composition and colonisation resistance can be influenced by host age, diet and sex^18^ ^19^, we next administered MEc30 to C57BL/6J mice under different host conditions (Figure 2A). Comparable fecal bacterial loads were observed between male and female mice, chow diet- and high-fat diet-fed mice, as well as young and aged mice at 30 days after administration (Figures 2B-D and Figures S3D-F), indicating that MEc30 colonisation is stable across multiple physiological contexts.

**Figure 2.**
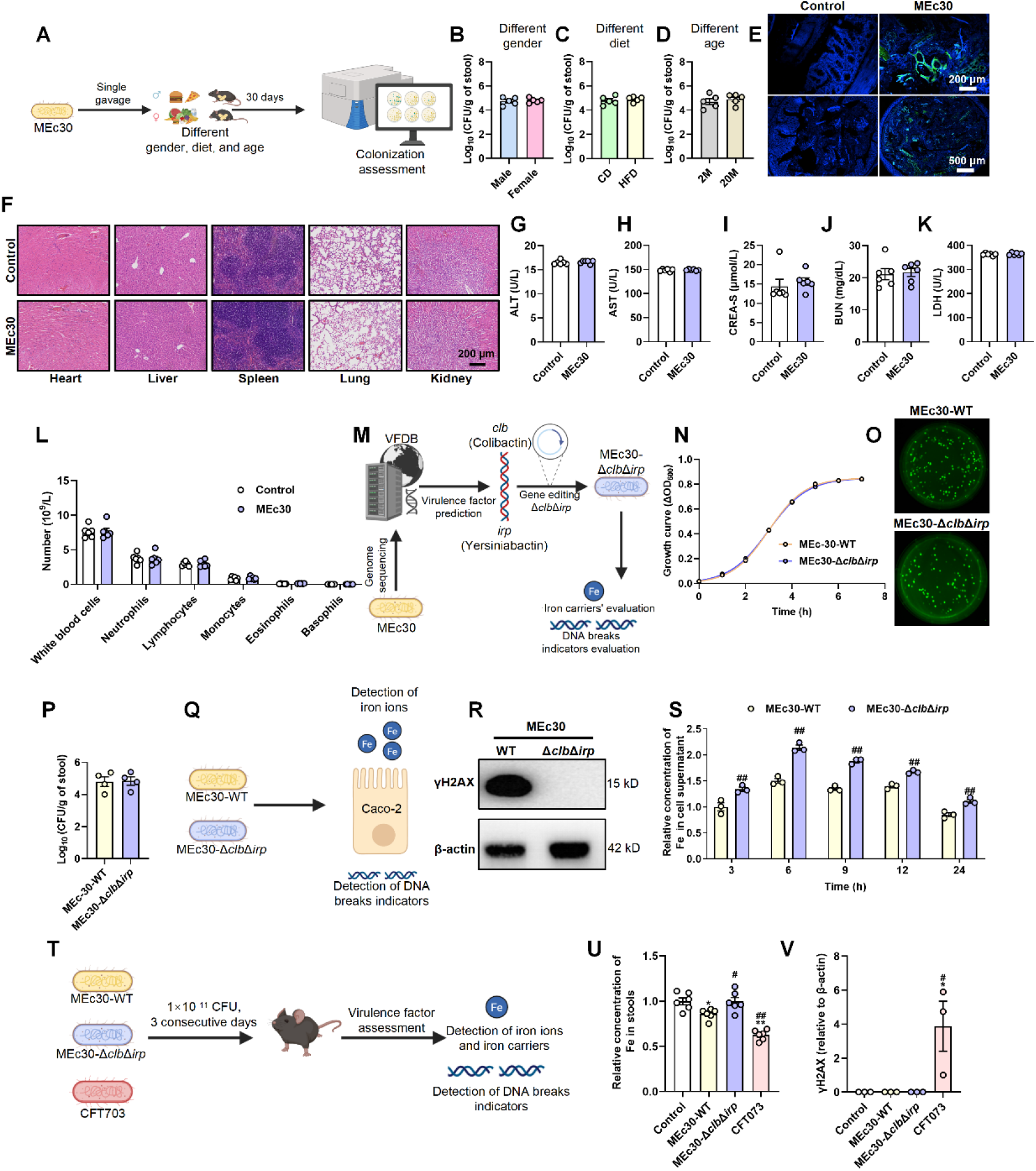
Evaluation of MEc30 colonisation robustness, biosafety, and genetic optimization *in vivo*. **(A)** Schematic overview of MEc30 colonisation assessment under different host conditions. Mice of varying sex (male and female), diet (chow diet (CD) and HFD), and age (2-month-old (2M) and 20-month-old (20M)) were orally administered MEc30 via a single gavage. Fecal samples were collected 30 days post-administration to evaluate intestinal colonisation levels. **(B-D)** Quantification of fecal CFU counts of MEc30 under different gender (male versus female) (B), diet (CD versus HFD) (C), and host physiological conditions-age (2M versus 20M) (D) at 30 days post-single gavage. **(E)** Representative FISH images showing MEc30 localization in the colon. Bottom panels show whole-section views; top panels display magnified images of the mucosal surface. Green fluorescence represents MEc30; blue indicates DAPI-stained nuclei. Scale bars: 500 μm (bottom), 200 μm (top). **(F)** Representative H&E-stained images of major organs (heart, liver, spleen, lung, kidney) from Control and MEc30-colonised mice. Scale bar, 200 μm. **(G-J)** Serum biochemical parameters, including alanine aminotransferase (ALT) (G), aspartate aminotransferase (AST) (H), creatinine (CREA-S) (I), blood urea nitrogen (BUN) (J), and lactate dehydrogenase (LDH) (K), in control and MEc30-treated mice. **(L)** Numbers of peripheral blood white blood cell (WBC) subpopulations (neutrophils, lymphocytes, monocytes, eosinophils, basophils). **(M)** Schematic overview of virulence factor identification and safety optimization of MEc30. Virulence-associated loci were predicted by VFDB analysis, followed by construction of the MEc30-Δ*clb*Δ*irp* derivative and evaluation of growth, colonization, and virulence-associated phenotypes. **(N)** Growth curve of MEc30-WT and MEc30-Δ*clb*Δ*irp* in LB medium. **(O)** Representative GFP-positive colony images of MEc30-WT and MEc30-Δ*clb*Δ*irp* recovered from fecal samples 30 days after a single oral administration. **(P)** Quantification of intestinal colonisation levels of MEc30-WT and MEc30-Δ*clb*Δ*irp*, shown as log_10_(CFU/g stool). **(Q)** Schematic illustration of the in vitro epithelial co-incubation assay used to evaluate iron-related phenotypes and DNA damage-associated indicators in Caco-2 cells exposed to MEc30-WT or MEc30-Δ*clb*Δ*irp*. **(R)** Western blot analysis of γH2AX and β-actin in Caco-2 cells after co-incubation with MEc30-WT or MEc30-Δ*clb*Δ*irp*. **(S)** Time-course quantification of Fe ion content in culture supernatants collected from Caco-2 cells co-incubated with MEc30-WT or MEc30-Δ*clb*Δ*irp*. **(T)** Schematic overview of the experimental design. Mice were orally administered PBS (Control) or 1×10^11^ CFU of the indicated *E. coli* strains (MEc30-WT, MEc30-Δ*clb*Δ*irp*, or CFT073) for three consecutive days, followed by *in vivo* assessment of iron-related phenotypes, including Fe ion and siderophore-associated changes, as well as DNA damage-associated indicators. **(U)** Relative concentration of Fe in stool samples from mice colonised with the indicated strains. **(V)** Densitometric quantification of colonic γH2AX protein levels relative to β-actin. All data are presented as mean ± SEM. The *p* values were determined by two-tailed Student’s t test in (B)-(S), The *p* values were determined by one-way ANOVA in (U) and (V). \**p* ≤ 0.05; \*\**p* ≤ 0.01 versus the Male/CD/2M respectively in (B)-(D), \**p* ≤ 0.05; \*\**p* ≤ 0.01 versus the Control group in (G)-(L), \**p* ≤ 0.05; \*\**p* ≤ 0.01 versus the Control group. #*p* ≤ 0.05; ##*p*≤ 0.01 versus MEc30-WT group in (N)-(V). *n* = 5 in each group in (B)-(D). *n* =4 in each group in (N)-(P). *n* = 3 in each group in (S) and (V). *n* = 6 in each group in (G)-(L) and (U).

We next assessed the spatial distribution of MEc30 along the intestine. MEc30 was detectable across multiple intestinal segments, including the duodenum, jejunum, ileum, cecum and colon, with enrichment in distal intestinal regions consistent with the known biogeography of gut microbial colonisation (Figures S3G-I)^20^. FISH analysis further showed that MEc30 signals were mainly located in the luminal and mucus-associated regions, rather than deep host tissue layers (Figure 2E and Figure S3J). This localization pattern is consistent with commensal *E. coli* behavior and distinct from pathogenic or pathobiont *E. coli* strains that can adhere to and invade epithelial or lamina propria compartments^21–23^.

We then evaluated whether stable MEc30 colonisation perturbed host physiology or the gut microbial ecosystem. At 30 days after a single administration, metagenomic analysis showed no appreciable changes in α-diversity or overall microbial community structure compared with control mice (Figures S4A-G). To further assess biosafety under supraphysiological exposure conditions^24^, mice were administered 1×10^11^ CFU MEc30 per day for three consecutive days and monitored for systemic toxicity (Figure S4H). MEc30-treated mice showed no significant body-weight changes, gross organ abnormalities or histological lesions in major organs, including the heart, liver, spleen, lung and kidney (Figure 2F and Figures S4I-M). Serum biochemical parameters, including alanine aminotransferase (ALT), aspartate aminotransferase (AST), serum creatinine (CREA-S), blood urea nitrogen (BUN), and lactate dehydrogenase (LDH), remained comparable between MEc30-treated and control mice (Figures 2G-L and Figures S5A-P). These results indicate that MEc30 does not detectably disturb host physiology, tissue integrity or gut microbiota composition under the tested conditions.

Although MEc30 showed a favorable *in vivo* safety profile, Virulence Factor Database (VFDB) analysis identified two putative virulence-associated loci, *clb* and *irp*, encoding colibactin- and yersiniabactin-associated functions, respectively (Figure 2M) ^24–27^. Because colibactin can induce DNA double-strand breaks and yersiniabactin mediates iron acquisition, we sought to further minimize potential safety risks for long-term therapeutic engineering. Guided by previous studies showing that deletion of the *pks*/*clb* island abolishes colibactin production and disruption of *irp1* reduces yersiniabactin biosynthesis^26^ ^28^, we finally generated MEc30-Δ*clb*Δ*irp* strain (Figures S6A-C). Deletion of these loci did not impair bacterial growth or 30-day intestinal colonisation capacity (Figures 2N-O). In Caco-2 epithelial cell co-incubation assays, MEc30-Δ*clb*Δ*irp* showed markedly reduced γH2AX accumulation compared with parental MEc30, a marker of DNA double-strand breaks and genotoxic stress^27^, indicating attenuation of colibactin-associated genotoxicity (Figures 2Q and 2R). In parallel, deletion of *irp* reduced the iron-scavenging phenotype, as reflected by altered iron ion levels in culture supernatants (Figure 2S).

Finally, we benchmarked MEc30 and MEc30-Δ*clb*Δ*irp* against the pathogenic *E. coli* strain CFT073, which contains both *clb* and *irp* loci and produces high levels of colibactin and yersiniabactin^25^ ^29^. Under high-dose administration, CFT073 showed stronger iron-scavenging activity, with lower extracellular or fecal iron levels and higher siderophore abundance, whereas MEc30-Δ*clb*Δ*irp* showed reduced siderophore production and restored iron levels compared with parental MEc30 (Figure 2U and Figures S6D-F). Moreover, CFT073 induced robust γH2AX accumulation in colon tissues, whereas MEc30 and MEc30-Δ*clb*Δ*irp* did not induce detectable intestinal γH2AX signals compared with controls (Figure 2V and Figure S6G). Together, these findings demonstrate that genetic removal of *clb* and *irp* further reduces potential virulence-associated risks without compromising bacterial fitness or colonisation capacity. Therefore, MEc30-Δ*clb*Δ*irp*, hereafter referred to as MEc30, was used as the safety-optimised native chassis for subsequent therapeutic engineering.

### Construction and characterization of MEc30-NA engineered bacteria

Nicotinic acid (NA), also known as niacin or vitamin B3, is a key precursor in the biosynthesis of nicotinamide adenine dinucleotide (NAD), an essential coenzyme involved in energy metabolism, DNA repair, and redox homeostasis^30^. Impaired NA/NAD metabolism has been associated with metabolic dysfunction, insulin resistance and ageing-related physiological decline^31^ ^32^. Although oral NA supplementation has long been used to improve lipid metabolism, its clinical application is limited by poor tolerance, short plasma half-life and peak-associated adverse effects^33^. We therefore sought to develop an engineered MEc30 strain capable of continuously producing NA within the gut.

To generate an NA-producing native chassis, we inserted *pyrZ*, which encodes a bifunctional enzyme involved in the conversion of quinolinic acid to nicotinic acid mononucleotide and NA^34^, into the chromosomal *attB* site of MEc30, generating MEc30-NA (Figure 3A and Figure S7A). This genomic modification did not impair bacterial growth compared with MEc30-WT (Figure 3B). MEc30-NA produced robust levels of NA in both LB culture and fecal culture conditions (Figure 3C and Figure S7B), indicating stable metabolic reprogramming of the native chassis.

**Figure 3.**
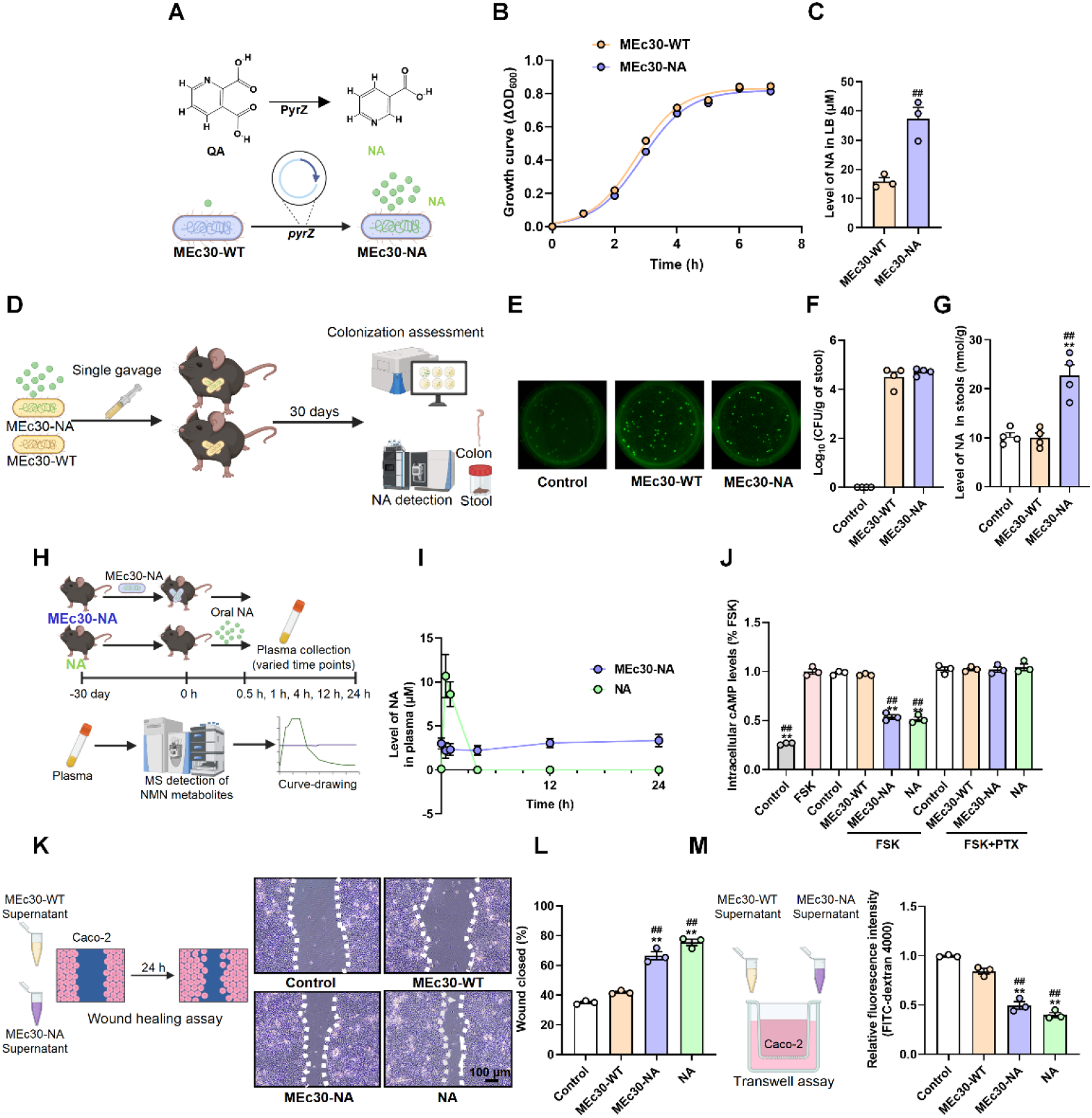
Construction and characterization of a MEc30-NA strain. **(A)** Schematic illustration of the construction of MEc30-NA. The *pyrZ* gene was introduced into MEc30 to reconstitute the QA-derived NA biosynthetic pathway, thereby enabling endogenous NA production. quinolinic acid (QA). **(B)** Growth curves of MEc30-WT and MEc30-NA in LB medium. **(C)** Quantification of NA abundance in LB culture supernatants of MEc30-WT and MEc30-NA strains, as determined by targeted liquid chromatography-tandem mass spectrometry (LC-MS/MS). **(D)** Schematic illustration of the in vivo colonisation and NA production assessment of MEc30-NA. Mice were orally administered a single dose of MEc30-WT or MEc30-NA and monitored for 30 days. Fecal and colonic samples were collected for colonisation quantification and NA detection. **(E-F)** Representative GFP-labeled colony images (E) and fecal CFU counts (F) in mice treated with PBS (Control) or colonised with MEc30-WT or MEc30-NA at day 30 post-gavage. **(G)** Quantification of NA in stools of mice treated with PBS (Control) or colonised with MEc30-WT or MEc30-NA at day 30 post-gavage. **(H)** Schematic illustration of the in vivo pharmacokinetic experiment. Mice in the MEc30-NA group were administered MEc30-NA and stable colonisation was confirmed. Then, mice in the NA group received a single oral dose of NA, while the MEc30-NA group received PBS as control. After that, blood samples were collected at 0, 0.5, 1, 4, 12, and 24 hours. Plasma concentrations of NA were measured by LC-MS/MS and plotted as concentration-time curves. **(I)** Relative abundance of NA in plasma at different time points following administration of MEc30-NA or NA. **(J)** Intracellular cAMP levels in Caco-2 cells indicating GPR109a activation. Cells were treated with forskolin (FSK) and bacterial supernatants from MEc30-WT, MEc30-NA, or NA, with or without pertussis toxin (PTX). cAMP was measured by ELISA. **(K)** Schematic design and representative images of wound-healing assays in Caco-2 cells treated with PBS (Control), MEc30-WT supernatants, MEc30-NA supernatants, or NA for 24 h. **(L)** Quantification of wound closure percentage in the corresponding groups. **(M)** Schematic and quantification of the FITC-dextran Transwell permeability assay used to assess epithelial barrier integrity. All data are presented as mean ± SEM. The *p* values were determined by two-tailed Student’s t test in (B)-(C) and (I); the *p* values were determined by one-way ANOVA in (G) and (J)-(M). \**p* ≤ 0.05; \*\**p* ≤ 0.01 versus the Control group. #*p* ≤ 0.05; ##*p*≤ 0.01 versus MEc30-WT group. *n* = 4 in each group in (B) and (F)-(I); *n*= 3 in each group in (C) and (J)-(M).

Following a single oral gavage, MEc30-NA and MEc30-WT achieved comparable intestinal colonisation levels at 30 days (Figures 3D-F). Stool metabolite analysis showed markedly increased fecal NA levels in MEc30-NA-colonised mice, and longitudinal monitoring confirmed sustained NA output throughout the 30-day observation period (Figure 3G and Figure S7C). Compared with direct oral NA administration, which caused a rapid plasma NA peak at approximately 0.5 h followed by decline to baseline within 24 h, MEc30-NA colonisation maintained a more stable elevation of plasma NA with substantially lower peak exposure (Figures 3H and 3I). These data suggest that MEc30-NA provides sustained intestinal NA production while avoiding the sharp systemic exposure peak associated with conventional NA administration.

Because NA is a ligand of GPR109a and NA-GPR109a signalling has been implicated in epithelial barrier protection and metabolic improvement^35^ ^36^, we next assessed whether bacterially derived NA could activate this pathway. MEc30-NA supernatants markedly suppressed forskolin-induced cAMP accumulation in Caco-2 cells, an effect comparable to free NA and reversed by pertussis toxin, indicating activation of canonical Gi-coupled GPR109a signalling (Figure 3J). MEc30-NA supernatants also increased the expression of *Gpr109a* and barrier-associated genes, including *Claudin1*, *Muc2*, *Occludin* and *Zo-1* (Figures S7D-H). Functionally, MEc30-NA promoted epithelial repair in wound-healing assays^37^ and reduced FITC-dextran flux across Caco-2 Transwell monolayers (Figures 3K-M), supporting enhanced epithelial restitution and barrier integrity. Together, these findings demonstrate that MEc30-NA continuously produces biologically active NA and activates epithelial GPR109a-associated barrier-protective signalling.

### MEc30-NA alleviated HFD-induced metabolic dysfunction

Oral administration of NA has been demonstrated to improve intestinal barrier permeability and alleviate HFD-induced metabolic dysfunction^36^. To evaluate whether endogenously produced NA from MEc30-NA exerts similar therapeutic effects, HFD-fed mice were treated with PBS, MEc30-WT, MEc30-NA or free NA after 8 weeks of HFD feeding, followed by an additional 5-week intervention (Figure 4A). At the endpoint, MEc30-NA maintained stable intestinal colonisation and markedly increased fecal NA levels, confirming sustained *in vivo* NA production (Figures S8A-C).

**Figure 4.**
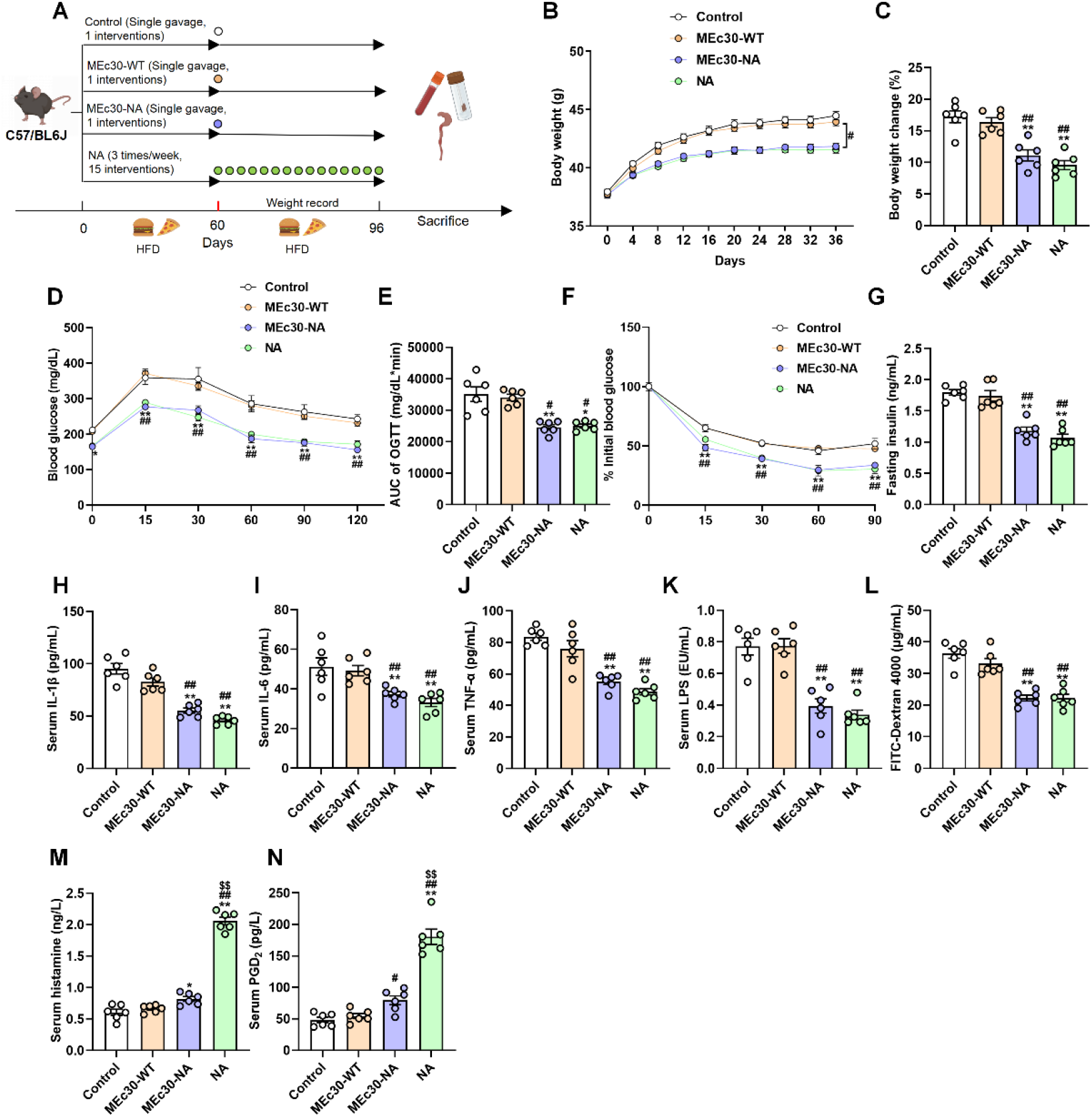
MEc30-NA alleviated HFD-induced metabolic dysfunction. **(A)** Experimental design. Mice were fed a high-fat diet (HFD) for 60 days and then orally administered PBS (Control), MEc30-WT, MEc30-NA, or nicotinic acid (NA) for an additional 36 days while continuing HFD feeding. **(B)** Body weight of mice recorded during the treatment period. **(C)** Percentage of body-weight change at the end of treatment. **(D-E)** Oral glucose tolerance test (OGTT), showing blood glucose levels over time (D) and the corresponding area under the curve (AUC). **(F)** Percentage of initial blood glucose during the insulin tolerance test (ITT). **(G)** Serum fasting insulin concentration. **(H-J)** Serum levels of IL-1β (H), IL-6 (I), and TNF-α (J). **(K)** Serum levels of LPS. **(L)** Serum FITC-dextran concentration 90 min after oral gavage to evaluate intestinal permeability. (**M-N**) Serum concentrations of histamine (N) and Prostaglandin D_2_ (PGD_2_). All data are presented as mean ± SEM. The *p* values were determined by one-way ANOVA in (B)-(N). \**p* ≤ 0.05; \*\**p* ≤ 0.01 versus the Control group. #*p* ≤ 0.05; ##*p*≤ 0.01 versus MEc30-WT group. *n*= 6 in each group.

Compared with HFD control and MEc30-WT groups, MEc30-NA significantly reduced body-weight gain and improved glucose metabolism, as shown by improved OGTT and ITT responses and reduced fasting insulin levels (Figures 4B-G). MEc30-NA also reduced liver weight, epididymal fat mass and circulating free fatty acid levels (Figures S8D-J), indicating broad improvement in HFD-induced metabolic dysfunction. MEc30-NA further reduced circulating LPS and pro-inflammatory cytokines, including IL-1β, IL-6 and TNF-α, and downregulated hepatic inflammatory gene expression (Figures 4H-K and Figure S8K). Consistent with the known role of NA-GPR109a signalling in intestinal barrier protection^36^, MEc30-NA upregulated colonic *Gpr109a* and barrier-associated genes, including *Claudin1*, *Muc2*, *Occludin* and *Zo-1*, increased colonic ZO-1 protein levels, and reduced FITC-dextran translocation *in vivo* (Figure 4L and Figure S8L-M). These results suggest that sustained microbial NA production improves metabolic homeostasis in part by suppressing metabolic inflammation and restoring intestinal barrier function.

Because pharmacological NA supplementation can induce flushing-related adverse responses mediated by prostaglandin D_2_ (PGD_2_) and histamine ^38^ ^39^, we next compared adverse-effect signalling between free NA and MEc30-NA. Direct NA administration markedly increased serum histamine and PGD₂ levels, whereas MEc30-NA induced only mild increases (Figures 4M and 4N), consistent with its lower systemic peak exposure. In addition, MEc30-NA was detectable in both proximal and distal colon and upregulated PPAR-α pathway-related genes, including *Ppara*, *Hmgcs2* and *Cbr3*, consistent with recent evidence that NA can promote barrier-protective colonic transcriptional remodeling through PPAR-α (Figures S8N-Q)^40^. Together, these findings demonstrate that MEc30-NA provides a sustained and better-tolerated microbial NA delivery strategy that improves HFD-induced obesity, insulin resistance, inflammation and intestinal barrier dysfunction.

### Construction and characterization of MEc30-mIL-10 engineered bacteria

IL-10 is a key anti-inflammatory cytokine that restrains excessive immune activation and maintains intestinal immune homeostasis^41^. Genetic variants in the IL10 locus are associated with increased risk of inflammatory bowel disease, and deficiency of IL-10 or IL-10 receptor signalling causes severe intestinal inflammation in both mice and humans^42–44^. Recent clinical studies of the oral IL-10 fusion protein AMT-101 have further supported the therapeutic potential of intestinal IL-10 supplementation in ulcerative colitis^45^ ^46^. However, repeated administration is required to maintain therapeutic exposure. We therefore reasoned that a durably colonizing engineered symbiont could provide a more sustained strategy for intestinal IL-10 delivery.

To generate an IL-10-producing native chassis, we engineered MEc30 to secrete murine IL-10 by chromosomally integrating an expression cassette containing the constitutive HCE promoter, the PelB secretion signal peptide and the murine *Il10* coding sequence into the *attB* site using a CRISPR-Cas9/λ-Red strategy (Figure 5A and Figure S9A)^47^. This genome-integrated design avoided plasmid-dependent expression and did not impair bacterial growth (Figure 5B). MEc30-mIL-10 secreted detectable IL-10 into culture supernatants during *in vitro* cultivation (Figures 5C and 5D).

**Figure 5.**
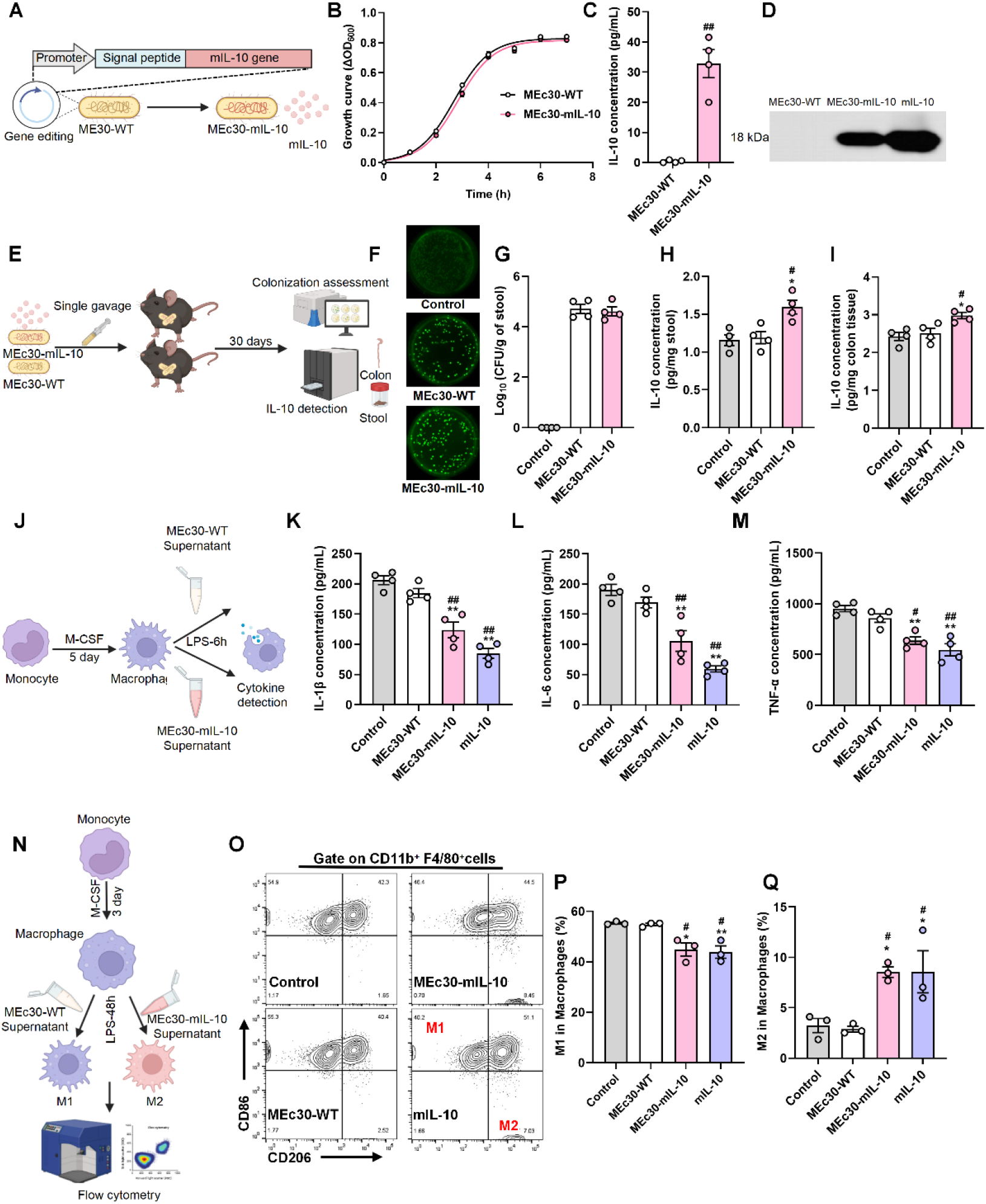
Evaluating the *in vitro* immunomodulatory activity of MEc30-mIL-10 strain. **(A)** Schematic illustration of the construction of the MEc30-mIL-10 strain. An expression cassette consisting of the constitutive HCE promoter, the PelB secretion signal peptide, and the murine IL-10 (*mIL-10*) coding sequence was integrated into the *attB* site of the MEc30 chromosome through CRISPR-assisted homologous recombination, enabling stable expression and secretion of mIL-10. **(B)** Growth curves of MEc30-WT and MEc30-mIL-10 in LB medium. **(C)** Quantification of IL-10 concentration in bacterial culture supernatants, measured by ELISA. **(D)** Western blot detection of IL-10 protein in culture supernatants of MEc30-WT, MEc30-mIL-10, and recombinant mIL-10. **(E)** Schematic illustration of the in vivo colonisation and IL-10 production assessment of MEc30-mIL-10. Mice were orally administered a single dose of MEc30-WT or MEc30-mIL-10 and monitored for 30 days. Fecal and colonic samples were collected for colonisation quantification and IL-10 detection. **(F)** Representative images of GFP-expressing colonies recovered from fecal samples of the indicated groups (Control, MEc30-WT, and MEc30-mIL-10) on LB agar plates at day 30 post-gavage, visualized under 488 nm fluorescence. **(G)** Quantification of fecal CFU in the indicated groups at day 30 post-gavage. **(H-I)** Quantification of IL-10 in stools (H) and colon tissue (I) of mice treated with PBS (Control) or colonised with MEc30-mIL-10 or MEc30-WT. **(J)** Schematic of *in vitro* experiment: BMDMs were differentiated with M-CSF for 5 days, stimulated with LPS, and treated with bacterial supernatants for cytokine detection. **(K-M)** Levels of IL-1β (K), IL-6 (L), and TNF-α (M) in BMDM culture supernatants measured by ELISA after treatment with Control, MEc30-WT, MEc30-mIL-10, or recombinant mIL-10. **(N)** Schematic of macrophage polarization assay: BMDMs were differentiated with M-CSF for 3 days, stimulated with LPS for 48 h in the presence of bacterial supernatants, and analyzed by flow cytometry. **(O)** Representative flow cytometry plots showing CD86 and CD206 expression gating on F4/80⁺CD11b⁺ macrophages. **(P-Q)** Percentages of M1 (P) and M2 (Q) macrophages among F4/80⁺CD11b⁺ cells as determined by flow cytometry. All data are presented as mean ± SEM. The *p* values were determined by two-tailed Student’s t test in (B)-(C). the *p* values were determined by one-way ANOVA in (H)-(Q). \**p* ≤ 0.05; \*\**p* ≤ 0.01 versus the Control group. #*p* ≤ 0.05; ##*p*≤ 0.01 versus MEc30-WT group. *n* = 4 in each group in (B)-(M). *n* = 3 in each group in (P)-(Q).

Following a single oral gavage, MEc30-mIL-10 and MEc30-WT achieved comparable intestinal colonisation levels at 30 days, indicating that the IL-10 expression cassette did not compromise colonisation fitness (Figures 5E-G). ELISA analysis showed significantly increased IL-10 levels in fecal supernatants and colonic tissues of MEc30-mIL-10-colonised mice compared with MEc30-WT controls (Figures 5H and 5I). Longitudinal monitoring further confirmed sustained fecal IL-10 production over 30 days (Figure S9B). These results demonstrate that MEc30-mIL-10 enables stable intestinal colonisation and continuous local IL-10 delivery.

We next assessed whether MEc30-derived IL-10 was biologically active. Because IL-10 suppresses M1 macrophage activation and promotes anti-inflammatory macrophage responses^48–50^, we treated LPS-induced bone marrow-derived macrophages with filtered bacterial supernatants. MEc30-mIL-10 supernatants significantly reduced the secretion of IL-1β, IL-6 and TNF-α compared with MEc30-WT supernatants (Figures 5J-M). Flow cytometry further showed that MEc30-mIL-10 supernatants inhibited M0 (CD11b^+^ F4/80^+^)-to-M1 macrophages phenotype (CD11b^+^ F4/80^+^ CD86^+^ CD206^-^) polarization and promoted M2 (CD11b^+^ F4/80^+^ CD86^-^ CD206^+^) polarization, as reflected by reduced M1 macrophages and increased M2 macrophages (Figures 5N-Q and Figures S9C-E). Together, these findings indicate that MEc30-mIL-10 secretes functional IL-10 capable of suppressing inflammatory macrophage activation and promoting anti-inflammatory macrophage polarization.

### MEc30-mIL-10 improves colitis caused by IL-10 gene defects

We next evaluated the therapeutic efficacy of MEc30-mIL-10 in *Il10*^-/-^ mice. At 16 weeks of age, *Il10*^-/-^ mice received a single oral dose of PBS, MEc30-WT or MEc30-mIL-10, while age-matched *Il10*^+/+^ mice served as healthy controls (Figure 6A). MEc30-mIL-10 stably colonised the intestine of *Il10*^-/-^ mice and significantly increased fecal IL-10 levels (Figures S10A-C). Compared with PBS(Control)- or MEc30-WT-treated *Il10*^-/-^ mice, MEc30-mIL-10 markedly alleviated body-weight loss, reduced colon shortening, decreased FITC-dextran translocation and restored colonic IL-10 levels (Figures 6B-E). Consistently, MEc30-mIL-10 reduced colonic IL-1β, IL-6 and TNF-α levels and ameliorated histological inflammation, indicating effective suppression of IL-10-deficiency-associated colitis (Figures 6F-J).

**Figure 6.**
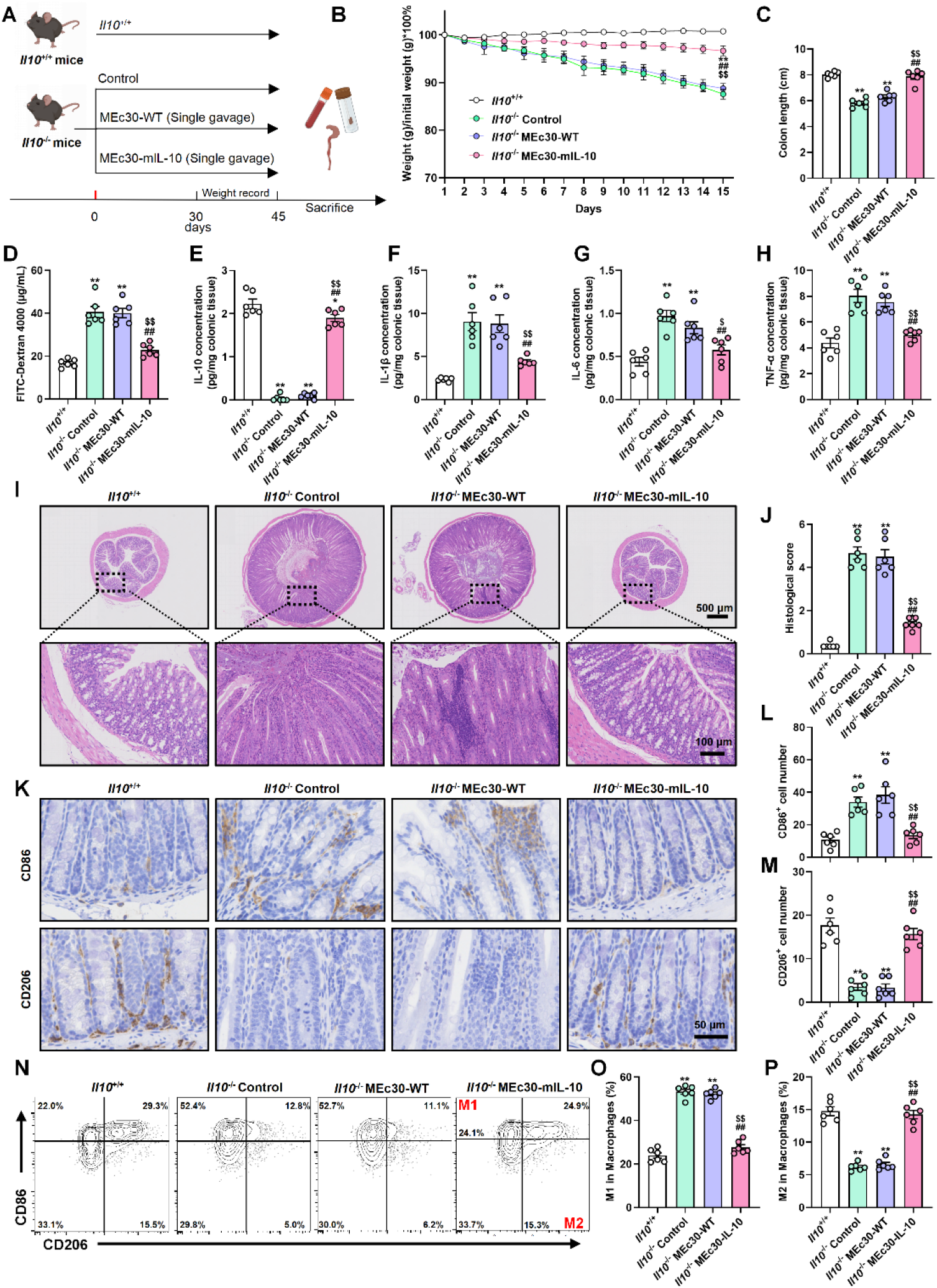
Evaluation of MEc30-mIL-10 in an IL-10-deficient mouse model of colitis and its effects on intestinal inflammation. **(A)** Schematic illustration of the experimental design. *Il10^-/-^* mice were orally administered a single dose of MEc30-WT, MEc30-mIL-10, or vehicle control and monitored for 45 days, with body weight recorded throughout the experiment. Age-matched *Il10^+/+^* littermates received PBS as healthy controls. **(B)** Body weight changes over 15 days in *Il10^-/-^* mice treated with PBS, MEc30-WT, or MEc30-mIL-10, and *Il10^+/+^* mice as a reference. **(C)** Colon length measurements at endpoint across treatment groups. **(D)** Serum FITC-dextran concentration 90 min after oral gavage to evaluate intestinal permeability. **(E–H)** Colonic tissue levels of IL-10 (G), IL-1β (H), IL-6 (I), and TNF-α (J) measured by ELISA. **(I)** Representative H&E staining of colon sections from each group (top: whole section; bottom: magnified region). Scale bars: 500 μm (top), 100 μm (bottom). **(J)** Histological scores based on inflammation, epithelial damage, and crypt architecture. **(K)** Representative IHC staining for CD86 and CD206 in colonic tissue sections from each group. Scale bar, 50 μm. **(L-M)** Quantification of CD86⁺ (L) and CD206⁺ (M) cell numbers per field in colonic tissue sections. **(N)** Representative flow cytometric contour plots showing colonic macrophage polarization in the indicated groups. Macrophages were identified as CD45^+^CD11b^+^F4/80^+^ cells and further classified into M1 (CD86^+^CD206^-^) and M2 (CD86^-^CD206^+^) subsets. **(O-P)** Quantification of the proportions of M1 macrophages (O) and M2 macrophages (P) in colonic tissues from the indicated groups. All data are presented as mean ± SEM. The *p* values were determined by one-way ANOVA in (B)-(M). \**p* ≤ 0.05; \*\**p* ≤ 0.01 versus *Il10*^+/+^ group. #*p* ≤ 0.05; ##*p*≤ 0.01 versus *Il10*^-/-^ Control group. $*p* ≤ 0.05; $$*p*≤ 0.01 versus *Il10*^-/-^ MEc30-WT group. *n*= 6 in each group.

We further examined macrophage responses in colonic tissues. Immunohistochemistry showed that MEc30-mIL-10 reduced pro-inflammatory M1 macrophage infiltration while increasing anti-inflammatory M2 macrophages in the colon (Figures 6K-M). Flow cytometry confirmed these findings, showing a decreased proportion of CD45^+^CD11b^+^F4/80^+^CD86^+^CD206^-^ M1 macrophages and an increased proportion of CD45^+^CD11b^+^F4/80^+^CD86^-^CD206^+^ M2 macrophages after MEc30-mIL-10 treatment (Figures 6N–P and Figures S10D-F). To further evaluate the therapeutic potential of MEc30-mIL-10 beyond the genetic *Il10*-deficient colitis model, we tested its efficacy in DSS-induced experimental colitis. MEc30-mIL-10 significantly attenuated body weight loss, reduced disease activity index, preserved colon length, alleviated histopathological injury, decreased MPO activity and pro-inflammatory cytokine expression, and improved intestinal barrier-related indicators compared with the DSS group (Figure S11). Together, these findings demonstrate that MEc30-mIL-10 restores local IL-10 availability, suppresses intestinal inflammatory responses, improves barrier integrity, and alleviates colitis in both genetic IL-10 deficiency and DSS-induced experimental colitis.

## DISCUSSION

Despite rapid progress in engineered bacterial therapeutics, achieving durable efficacy, predictable colonisation and acceptable biosafety remain a major translational challenge. Several engineered strains have shown promising preclinical activity, but their therapeutic effects may be limited when sustained molecular delivery is required. For example, SYNB1618 and SYNB1934 were developed for phenylketonuria and showed phenylalanine-lowering activity in early studies, but later clinical development did not achieve the expected endpoints^51^. Similarly, engineered *Lactococcus lactis* secreting IL-10 reduced colitis in mice and showed safety in early clinical testing, but subsequent clinical efficacy was insufficient^52–54^. One potential limitation of such strategies is their transient intestinal persistence, requiring repeated administration and potentially limiting sustained therapeutic output^55^ ^56^. These examples suggest that durable host colonisation may be particularly important for engineered bacterial therapies designed to provide long-term local delivery of anti-inflammatory or metabolic molecules.

EcN has been widely used as a chassis for oral engineered bacterial therapies because of its genetic tractability and probiotic history. However, its colonisation capacity is often variable across hosts and experimental settings, and strategies to improve persistence frequently require antibiotic preconditioning or auxiliary delivery system^57^. Encapsulation or hydrogel-based approaches can improve bacterial retention and targeted delivery, but they may introduce additional complexity, variable release kinetics and material-associated safety concerns^58^ ^59^. In our study, EcN showed only transient intestinal persistence after oral administration, whereas MEc30 achieved stable long-term colonisation without antibiotic pretreatment or auxiliary delivery systems. This feature directly addresses a central limitation of current live bacterial therapeutics: the difficulty of maintaining sufficient in vivo abundance for sustained therapeutic activity.

Recent studies have highlighted the potential of host-derived native *E. coli* strains as chassis for persistent intestinal transgene delivery^6^ ^8^. However, whether durable engraftment is a general property of native isolates has remained unclear. Using conventional fecal isolation, we initially obtained 10 native *E. coli* strains, but none maintained detectable colonisation beyond 30 days. We therefore applied an *in situ* cultivation strategy and screened 116 additional native isolates, from which MEc30 emerged as a rare strain capable of lifelong intestinal persistence. One possible explanation is that *in situ* cultivation may reduce *ex vivo* growth bias and better preserve host-adapted bacteria with durable niche-occupation capacity^17^. These findings extend prior work on native bacterial chassis by showing that durable engraftment is a highly strain-specific trait that requires systematic screening rather than simple selection of host-derived isolates.

Safety is another essential consideration for long-term live biotherapeutic development. Although MEc30 did not detectably disturb host physiology, tissue integrity or gut microbial community structure under the conditions tested, genomic screening identified the putative virulence-associated *clb* and *irp* loci. Because colibactin and yersiniabactin have been linked to genotoxicity and iron-scavenging-associated pathogenic traits^25^ ^26^, we deleted these loci to generate a safety-optimised derivative. Importantly, *clb/irp* deletion reduced colibactin- and yersiniabactin-associated activities without impairing bacterial growth or colonisation capacity. This step provides a framework for balancing native colonisation fitness with rational biosafety optimization, which will be important for future clinical translation of engineered native symbionts.

Using this safety-optimised chassis, we demonstrated two therapeutic applications relevant to chronic intestinal inflammatory and metabolic diseases. MEc30-mIL-10 restored intestinal IL-10 availability, suppressed inflammatory macrophage activation, promoted anti-inflammatory macrophage polarization and alleviated colitis in *Il10*^-/-^ mice. This supports the feasibility of using durably colonizing bacteria for sustained local cytokine delivery in host-deficiency-associated intestinal inflammation. In parallel, MEc30-NA provided continuous intestinal nicotinic acid production, activated epithelial GPR109a-associated barrier signalling, improved insulin sensitivity, reduced metabolic inflammation and avoided the sharp systemic peak exposure associated with conventional NA administration. Beyond GPR109a, MEc30-NA was detected in both proximal and distal colonic contents and upregulated PPAR-α pathway-related proximal colonic identity genes in the colon, supporting PPAR-α-associated barrier-protective colonic transcriptional remodeling. Its potential application in other metabolic or barrier-associated diseases requires disease-specific validation. Notably, MEc30-WT showed a modest trend toward improving epithelial barrier function, consistent with the reported barrier-supporting effects of commensal or probiotic bacteria^60^ ^61^, whereas MEc30-NA produced a stronger protective effect. Together, these findings show that MEc30 can serve as a modular native chassis for both therapeutic protein delivery and metabolic molecule production.

Although MEc30 exhibited stable colonization, genetic stability, and an improved biosafety profile in mice, several limitations should be noted. Our study was conducted in mouse-derived native *E. coli* and murine disease models, and thus its translational potential in the human intestinal ecosystem remains to be established. In addition, although the engineered platform successfully rescued IL-10 deficiency-associated colitis, the genetic-disease application was tested in only one host-deficiency model. Whether this strategy can be extended to other genetic deficiency settings requires further investigation. Furthermore, the molecular and ecological determinants underlying the exceptional lifelong colonisation of MEc30 remain incompletely understood, limiting mechanistic insight into this durable engraftment phenotype. Moreover, although our data support durable colonisation and sustained functional output in mice, longer-term follow-up in additional disease contexts and further mechanistic dissection will be important for future translational development. Future translation will also require the identification of human-compatible durable colonisers, evaluation of host specificity and horizontal gene transfer risks, development of controllable clearance or kill-switch systems, and resolution of manufacturing and regulatory challenges associated with live biotherapeutic products. Evaluation in more clinically relevant systems will therefore be essential to establish the long-term safety, stability, and therapeutic robustness of this platform.

## Lead Contact

Further information and requests for resources and reagents should be directed to and will be fulfilled by the lead contact, Kai Wang.

## Data availability statement

All data supporting the findings of this study are available from the Lead Contact upon reasonable request. This study does not include original code. Any additional information required to reanalyze the data is also available from the Lead Contact upon request. The bacterial strains are available directly from the authors upon reasonable request.

## Contributors

C. J., K. W. conceptualized and designed the study. Q. Z., Y. D., S. Y., H. M., Y. W., S. G., and Y. P., performed the experiments and analyzed the data. C. J., and K. W. supervised the study. Q. Z., Y. D. and C. J. wrote the manuscript. Q. Z. and Y. D. contributed equally to this work. All authors edited the manuscript and approved the final manuscript.

## Ethics approval

All animal experiments were approved by the Institutional Animal Care and Use Committee of Peking University Health Science Center (approval NO. BCJB0012) and were performed in accordance with the Guide for the Care and Use of Laboratory Animals.

