## Supplementary material for "A durably colonizing engineered native symbiont enables sustained intestinal delivery for metabolic dysfunction and colitis": Method

**METHODS**

**Mice and Treatments**

All animal protocols were approved by the Institutional Animal Care and Use Committee (IACUC) of Peking University Health Science Center (permit: BCJB0012). C57BL/6J mice (GemPharmatech, N000192) and C57BL/6JGpt-Il10$^{em1Cd4885}$/Gpt mice (*Il10*$^{-/-}$mice) (GemPharmatech, T005959) were purchased from GemPharmatech. All mice had access to food and water ad libitum and were kept under a strict 12-h light/dark cycle at a controlled temperature (23 ± 2°C).

Age-matched (8-week-old) male mice were used in this study and were randomly assigned to experimental groups using a random-number generator after acclimatization. A double-blind design was implemented whenever feasible. Group allocation codes were generated and maintained by an investigator who was not involved in animal handling, outcome assessment or data analysis. Treatment preparations and biological samples were labelled using coded identifiers, and investigators responsible for animal handling, treatment administration, sample collection, outcome assessment and statistical analysis were blinded to group allocation throughout the study. Group identities were not revealed until all data collection, outcome assessments and primary statistical analyses had been completed. Body weights of mice across groups were not significantly different before treatments were applied. Mice were euthanized by $CO_2$ asphyxiation, and no mice were excluded from the analysis. Mice were gavaged with 200 μL of either PBS or ~$1\times10^{10}$ CFU/mL of the corresponding bacterial strain unless otherwise stated. For all colonization experiments, mice were housed individually (one mouse per cage) throughout the study period to minimize cage effects, cross-transmission, and environmental reseeding. Mice from different experimental groups were maintained under strictly separated housing conditions, and animal handling as well as fecal sample collection were performed individually to avoid cross-contamination between cages.

In order to evaluate the colonization of the intestinal tract of mice with proresident bacteria, the 8-week-old C57BL/6J male SPF mice were administrated once with 12 strains including EcN, MEc3, MEc4, etc. Intestinal colonization of the above proresident bacteria was detected at 1, 3, 7, 14, 30, 60, 120, 360, and 720 days until colonization of the strain was not detectable in all mice in the group.

To explore the optimal colonization dose of MEc30, 8-week-old C57BL/6J male SPF mice were given $1\times10^4$ CFU/mL, $1\times10^6$ CFU/mL, $1\times10^8$ CFU/mL, and $1\times10^{10}$ CFU/mL MEc30. Fecal samples were collected at day 1 and day 30 to detect MEc30 colonization. During the entire colonization-monitoring period, each mouse was maintained in a separate cage to prevent microbial transmission between animals and to ensure that long-term persistence reflected true colonization rather than re-exposure from the environment or cage mates.

To detect the colonization of MEc30 in mice with different diets, ages and genders. We administered MEc30 to female and male mice, mice aged 2 months and 20 months, mice on the CD diet and mice on the HFD diet (Research Diets, D12492). Fecal samples were collected at day 1 and day 30 to detect MEc30 colonization. Mice in this experiment were also housed individually in separate cages throughout the study.

To investigate the efficacy of MEc30-mIL-10 in the treatment of colitis caused by genetic defects in IL-10, we administrated MEc30-mIL-10, MEc30-WT and PBS controls in 4-month-old *Il10*$^{-/-}$ mice. At the age of 5 months, the weight of mice began to decrease, and the weight change of mice in each group was detected. At the same time, the mice were killed after 15 days of weight measurement, and the colonization of MEc30-mIL-10 and MEc30-WT in feces was detected.

To assess the metabolic effects of MEc30-NA, 8-week-old male C57BL/6J mice were fed an HFD for 8 weeks. After that, mice were treated with (1) PBS, (2) MEc30-WT (single oral dose), (3) MEc30-NA (single oral dose), or (4) nicotinic acid (NA; MCE, HY-B0143) at 150 mg/kg, three times per week. At study endpoint, feces, blood, liver, and colon tissue were harvested for analysis of colonization and host metabolic parameters.

For all in vivo experiments, the individual mouse, rather than the cage, was considered the experimental unit (*n*).

**Bacterial strains and culture conditions**

Native *E. coli* strains were isolated from murine fecal samples and cultured in LB broth or on LB agar plates at 37°C under aerobic conditions unless otherwise indicated. The probiotic strain *Escherichia coli* Nissle 1917 (EcN; BNCC 361741, BeNa Culture Collection, China) was used as a reference strain, and uropathogenic *E. coli* CFT073 (BNCC 354053, BeNa Culture Collection, China) was used as a control in selected virulence-related assays. BNCC is a commercial microbial strain provider, and both strains are publicly accessible and commercially available. A total of 116 native isolates (designated MEc1-MEc116) were obtained in this study.

For antibiotic selection, LB medium was supplemented with kanamycin (Sigma-Aldrich, D403; 50

μg/mL), spectinomycin (Sigma-Aldrich, S0692; 50–100 μg/mL as indicated), ampicillin (Sigma-Aldrich, BP021), gentamicin sulfate (Sigma-Aldrich, E003632), streptomycin (Sigma-Aldrich, S9137), or chloramphenicol (Sigma-Aldrich, C1919) when appropriate. Anhydrotetracycline (Aladdin, A276282) was used when required for gene-editing procedures.

For genetic engineering, MEc30-GFP was used as the chassis strain. Construction of MEc30-mIL-10 and MEc30-NA was performed via a CRISPR-Cas9 system mediated by the Bimonder editing platform (Bionedit), followed by plasmid curing and sequence validation. Engineered strains were routinely maintained in antibiotic-free LB after confirmation. The codon-optimized *pyrZ* sequence and murine *Il10* coding sequence used to construct MEc30-NA and MEc30-mIL-10, respectively, are provided in Supplementary Materials 3, Table 1.

**Cell culture conditions**

Caco-2 cells (iCell, h032) were cultured in DMEM medium (Gibco, C11995500BT) supplemented with 10% fetal bovine serum and 1% penicillin/streptomycin at 37°C in a humidified incubator with 5% $CO_2$.

Bone marrow-derived macrophages (BMDMs) were maintained at 37°C in a humidified atmosphere with 5% $CO_2$ and were confirmed to be mycoplasma-free. For BMDM differentiation, RPMI-1640 medium (Gibco, C11995500BT) supplemented with 10% fetal bovine serum, 1% penicillin/streptomycin, and recombinant mouse M-CSF (BioLegend, 576404) was used.

**Isolation and in situ enrichment of native intestinal *E. coli***

For the initial isolation of native *E. coli* strains, five C57BL/6J mice from different cage positions were randomly selected. Fresh fecal samples were collected from each mouse and processed individually. Feces were suspended in sterile PBS (1:10, w/v), homogenized, serially diluted, and plated directly onto lactose-containing MacConkey agar medium (Solarbio, M8560). After incubation, multiple colonies showing typical *E. coli*-like morphology and lactose-fermenting phenotype (pink colonies) were manually picked from each mouse by visual inspection and purified by restreaking. Candidate isolates were then subjected to 16S rRNA gene PCR and sequencing for taxonomic confirmation. Based on the sequencing-confirmed isolates, two native *E. coli* strains were ultimately retained from each mouse, yielding a total of 10 original native *E. coli* isolates.

To further enrich and recover native *E. coli* strains with better preservation of their ecological competitiveness, we subsequently adapted an in situ cultivation approach using a diffusion chamber design as described previously[1]. Fecal samples were collected from C57BL/6J mice in different facilities, genders, diets, and ages. Fresh feces were suspended in sterile PBS (1:10, w/v) and homogenized. Approximately 1 mL of fecal suspension was mixed with agar and sealed between two 0.22-μm membrane filters (Biodee, DE-NKY-0456) using sterile stainless-steel washers to form diffusion chambers. The chambers were incubated within freshly collected fecal material under anaerobic conditions for 3 days, allowing nutrient and signaling exchange while retaining the bacterial cells inside.

After incubation, chamber contents were recovered, serially diluted, and plated on lactose-containing MacConkey agar. Colonies were selected manually by visual inspection rather than by an automated algorithm. Specifically, multiple colonies showing typical *E. coli*-like morphology and lactose fermentation (pink color) were picked from the plates and purified by restreaking. Candidate isolates were subsequently confirmed by 16S rRNA gene PCR and sequencing, and a total of 116 sequencing-validated native *E. coli* isolates were obtained for subsequent analyses. Phylogenetic identification was performed by BLAST analysis against the NCBI 16S rRNA database.

**Growth curves of tested strains**

Fresh strains ($2\times10^7$ CFU/mL) at 2 μL were inoculated into freshly prepared medium (200 μL), then growth anaerobically for 8 h. The bacterial growth kinetics were analyzed using a Spectra Max 190 microplate reader (Molecular Devices Inc.) by measuring the optical density ($OD_{600}$).

**Native *E. coli* antibiotic resistance testing**

Antibiotic resistance of the native *E. coli* was referred to the study by Ma et al[2]. After culturing the test *E. coli* to the logarithmic growth phase, the bacterial culture is diluted to a concentration of $1\times10^8$ CFU/mL using sterile PBS. Then, different strains are treated with working concentrations of common antibiotics and incubated at 37°C for 12-16 hours. The $OD_{600}$ values of the cultures in each group are measured using a spectrophotometer and compared to the $OD_{600}$ value of the control group (without antibiotics) to evaluate the bacterial sensitivity to the antibiotics. We define a reduction of over 65% in the $OD_{600}$ value compared to the control group as significant inhibition of bacterial growth, indicating sensitive; otherwise, it is considered resistance.

**GFP-Fluorescent colony count**

After incubating a plate coated with bacterial solution at 37°C overnight (12-16 hours), GFP positive colonies were detected in the ChemiDoc imaging system controlled by Image Lab software (Bio-Rad, 12003153) under 488 excitation light. ImagePro Plus software was used to further quantify the number of GFP-positive colonies in the images.

**Colonization Assessment**

For feces, we collected 2-4 fecal pellets from each mouse and weighed them. Add 10 μL sterile PBS per mg sample, homogenize feces, centrifuge supernatant, and dilute the diluted bacterial solution according to the appropriate ratio and multiple ratios, and apply the diluted bacterial solution on the LB plate to calculate CFU/g feces.

For tissues, mice were euthanized and tissue samples were harvested and immediately placed on ice. Dissection tools were disinfected between animals. Samples were processed and plated as described for stool. Collected and tested tissues include duodenum defined as the first 10 cm of small intestine, jejunum defined as the next 10 cm of small intestine, Ileum defined as the next 5 cm of jejunum, cecum, and colon.

**FISH fluorescence in situ hybridization**

Colon tissues were fixed in 4% paraformaldehyde in PBS, embedded in paraffin, and sectioned. Sections were pretreated with the FISH probe reaction buffer kit (Genepharma, F26501/100) and then hybridized with the probes. The probes were MEc30-GFP: 5′FAM- TGTCTTGTAGTTCCCGTCATC. Sections were incubated in 2 μM FISH probes (Qing Ke Zi Xi Biotechnology Co) for 14 h at 37 °C in a humidified chamber. After that, sections were then incubated in the dark with DAPI for 10 min at room temperature. Images were analyzed using a Nikon NSPARC confocal microscope controlled by NIS-Elements viewer (version 5.12).

**Metagenomic analysis of fecal microbial composition**

After 15 days of MEc30 colonization, fresh fecal samples were collected from control and MEc30-treated mice and stored at −80°C. Microbial genomic DNA was extracted for metagenomic library construction. Sequencing libraries were prepared using the Ultra DNA Library Prep Kit for Illumina according to the manufacturer's instructions. DNA concentration was measured using a Qubit DNA Assay Kit on a Qubit 2.0 fluorometer, and library quality and insert size were assessed using an Agilent Bioanalyzer 2100. Libraries with qualified concentrations were clustered on a cBot Cluster Generation System and sequenced on an Illumina HiSeq 4000 platform to generate 150-bp paired-end reads.

Raw metagenomic reads were processed using bioBakery tools. Low-quality reads and adapter sequences were removed, and host-derived reads were filtered using KneadData with integrated Bowtie2. Taxonomic profiling was performed using MetaPhlAn with default parameters. Alpha diversity, including observed species richness and Shannon index, was calculated based on taxonomic profiles. Beta diversity was assessed by principal coordinates analysis (PCoA) based on community composition distance matrices.

**Hematological and serum biochemical analysis**

After 15 days of MEc30 colonization, serum and whole blood were collected from both control and MEc30-treated mice. Samples were submitted to the Laboratory Animal Science Department at Peking University Health Science Center for hematological and biochemical analysis.

**H＆E analysis**

For histological analysis of colonic tissue, proximal colon segments were immediately removed and fixed in 4% paraformaldehyde in PBS for 24 h at room temperature. Post 10% formalin, the colon samples were routinely dehydrated, and then embedded in paraffin and thin sections (5 μm) were cut and deposited on glass slides. The paraffin sections were stained with H&E.

To assess the toxicological effects of MEc30 on major organs, the major organs of mice in the Control group and the MEc30 group: heart, liver, spleen, lung and kidney, were treated according to the intestinal H&E method described above.

**In vitro infection for DNA damage analysis**

Caco-2 cells were infected with MEc30-WT, MEc30-Δclb, or MEc30-ΔclbΔirp at an MOI of 100. To maximize bacterial–epithelial contact, plates were centrifuged at 300 × g for 1 min and incubated at 37°C with 5% $CO_2$ for 6 h. Cells were then washed three times with PBS and cultured in complete medium containing gentamicin sulfate (100 μg/mL) for 30 min, followed by incubation in complete medium containing gentamicin sulfate (50 μg/mL) until 24 h after infection initiation. Cells were then harvested for analysis of DNA damage marker γH2AX using anti-phospho-H2AX antibody (ABclonal, AP0099).

**In vitro infection for supernatant iron measurement**

Caco-2 cells were infected with MEc30-WT, MEc30-Δ*clb*, or MEc30-Δ*clb*Δ*irp*, and an uninfected group was included as the control. Cells were cultured in 6-well plates to 70–80% confluence before infection. The indicated bacterial strains were added to Caco-2 cells at the designated multiplicity of infection (MOI of 100), and the plates were incubated at 37 °C with 5% $CO_2$. At 3, 6, 9, 12, and 24 h after infection[3], culture supernatants were collected and centrifuged at 3,000 rpm for 5 min to remove cells and debris. The clarified supernatants were then used for subsequent determination of iron ion content according to the manufacturer's instructions.

**Iron quantification in bacterial cultures and fecal samples**

Ferrous ion concentrations in bacterial supernatants and mouse samples were determined using a ferrozine-based Iron Kit (BIOSINO, 100020290) according to the manufacturer's instructions.

For in vitro analysis, *E. coli* strains were cultured in LB medium (for total Fe measurement) to mid-log phase ($OD_{600}$=0.6). Cultures were centrifuged and supernatants were filtered through 0.22 μm membranes. To determine total iron, aliquots were incubated with freshly prepared ascorbic acid to reduce $Fe^{3+}$ to $Fe^{2+}$, followed by reaction with ferrozine reagent. Absorbance was measured at 570 nm using a microplate reader. Iron concentrations were normalized to culture density ($OD_{600}$).

For in vivo analysis, fecal samples were collected from mice after treatment. Fresh stools (100 mg) were homogenized in 1 mL PBS. After cooling and centrifugation, supernatants were neutralized to pH 4-5 using NaOH and subjected to ferrozine assay as above. The optical density was read at 570 nm, and total Fe concentration was determined against $FeSO_4$ standards.

**Siderophore quantification in bacterial cultures and fecal samples**

Siderophore concentrations were measured using a CAS siderophore detection kit (Qiyunbio, QM5114) according to the manufacturer's instructions.

For *in vitro* assays, *E. coli* strains were cultured in M9 medium (Sangon, A507024-0250) for 3 days. Supernatants were collected after centrifugation and filtration. Siderophore production was assessed using the Chrome Azurol S (CAS) liquid assay, in which bacterial supernatants were mixed 1:1 with CAS reagent and incubated for 30 min at room temperature. Absorbance was measured at 630 nm, and siderophore activity was expressed as the percentage of siderophore units (%SU) using the formula: $\%SU = (A_r - A_s)/ A_r \times 100$, where $A_r$ and $A_s$ represent the absorbance of the reference and sample, respectively.

For in vivo assays, fecal samples (100 mg) were homogenized in 1 mL PBS. After cooling and centrifugation, supernatants were neutralized to pH 4-5 using NaOH and subjected to the same ferrozine and CAS assays to determine siderophore levels, respectively. The siderophore activity of each strain in in vitro assays was normalized to that of CFT073, which was set as 1. For in vivo experiments, siderophore levels in fecal samples were normalized to the Control group, which was set as 1.

**Western blot**

Colon tissues were homogenized in RIPA buffer containing protease and phosphatase inhibitors, and total proteins were extracted. Caco-2 cells were lysed in the same buffer. Culture supernatants from MEc30-WT and MEc30-mIL-10 were also collected for protein analysis. Samples were resolved by SDS-PAGE and transferred to PVDF membranes. Membranes were blocked with 5% skim milk for 1 h at room temperature and incubated with primary antibodies, including anti-IL-10 (Abcam, ab310329), anti-β-actin (ABclonal, AC026), and anti-phospho-H2AX (ABclonal, AP0099), followed by HRP-conjugated goat anti-rabbit IgG (ABclonal, AS014). Blots were visualized using chemiluminescence and quantified with a ChemiDoc imaging system controlled by Image Lab software.

**ELISA**

For analysis of inflammatory cytokines in BMDM polarization experiments, culture supernatants were collected and assayed using mouse IL-1β, IL-6, and TNF-α ELISA kits (Acmec; AC16536, AC16541, and AC16568, respectively).

For detection of IL-10 secreted by MEc30-mIL-10, bacterial supernatants were collected by centrifugation and assayed using a mouse IL-10 ELISA kit (Acmec, AC16544).

For fecal IL-10 detection, fecal samples were first weighed, then resuspended in PBS, homogenized, and centrifuged; the supernatant was collected and analyzed for IL-10 concentration using the ELISA kit.

For the detection of IL-10, IL-1β, IL-6, TNF-α in colon tissue, the colon tissue was weighed and ground under liquid nitrogen, and a specific volume of PBS resuspension tissue containing PMSF was added according to the ELISA kit instructions. For the detection of ZO-1 in colon tissue, the colon tissue was weighed, homogenized in PBS, subjected to two freeze–thaw cycles, and centrifuged. The supernatant was collected, and ZO-1 levels were quantified using a mouse ZO-1 ELISA kit according to the manufacturer’s instructions. For the detection of circulating IL-10, IL-1β, IL-6, and TNF-α, blood was collected and serum was isolated for cytokine quantification. After centrifugation, the concentration of IL-1β, IL-6, TNF-α in the tissue was detected by ELISA.

For serum insulin quantification, blood was collected after a 6-h fast, and insulin was measured using a mouse insulin ELISA kit (Cusabio, CSB-E05071m). The resulting values were used to calculate HOMA-IR.

For systemic histamine and $PGD_2$ measurement, serum histamine and $PGD_2$ were quantified using mouse histamine and prostaglandin $D_2$ ELISA kits (Elabscience, E-EL-0032 and E-EL-0066).

**Plasma endotoxin measurement**

Plasma endotoxin levels were determined using an endpoint chromogenic Limulus amebocyte lysate (LAL) assay kit (Yeasen Biotechnology, Shanghai, China) according to the manufacturer’s protocol. Briefly, 50 μL of plasma or endotoxin standard was transferred into pyrogen-free tubes and mixed with 50 μL of LAL reagent. After incubation at 37°C for 8 min, 50 μL of chromogenic substrate was added, followed by a further incubation at 37°C for 6 min. The reaction was terminated by adding 250 μL of stop solution. After standing at room temperature for 5 min, absorbance was measured at 545 nm. Plasma endotoxin concentrations were calculated from the standard curve and expressed as EU/mL.

**Biochemical analysis**

Total cholesterol (TC), triglycerides (TG), and free fatty acids (FFAs) were measured using commercially available kits, including TC kit (BIOSINO, 100060092), TG kit (BIOSINO, 100000220), and FFAs kit (Beyotime, S0215S), according to the manufacturers' instructions.

**Isolation of mouse monocytes and induction of BMDMs**

Isolation and induction of mouse monocytes refer to previously published articles[4]. In short, mouse BM-derived macrophages were isolated from C57BL/6J mice. The bone marrow was rinsed into the medium (RPMI 1640, Invitrogen, 31870082), filtered through a 70 μm sieve, and then centrifuged. Cell density was adjusted to $1\times10^6$/mL and cultured in macrophage medium (RPMI 1640) supplemented with 10% fetal bovine serum, 1% penicillin/streptomycin, recombinant mouse M-CSF (BioLegend, 576404; 1 ng/mL), for 5 days at 37 °C in an incubator containing 5% $CO_2$. After culture, flow cytometry was used to identify the induced differentiation results and follow-up experiments were carried out. After that, BMDMs was activated with MEc30-mIL-10 engineered bacteria supernatant or control in the presence of LPS (Solarbio, L8880; 100 ng/mL) for 6 hours. After culture, the cell supernatant was collected to detect the expression of inflammatory factors. For the macrophage polarization experiment, monocytes were cultured for 3 days after the addition of M-CSF to differentiate into M0 cells. Then, the supernatant filtrate of MEc30-mIL-10 engineered bacteria and control bacteria and recombinant mouse IL-10 (BioLegend, 575802) was added to the medium containing 100 ng/mL LPS. After 2 days of continuous culture, the cells were digested with pancreatic enzymes and the macrophage polarization was detected by flow cytometry.

**Flow cytometry**

For flow cytometric analysis of colonic macrophages, fresh colon tissues were collected from mice, opened longitudinally, washed thoroughly with cold PBS to remove luminal contents, and cut into small pieces. To remove epithelial cells, tissues were incubated in HBSS containing 5 mM EDTA, 1 mM DTT, and 3% FBS at 37℃ with shaking for 20 mins. The remaining tissues were then digested in RPMI-1640 containing collagenase IV, neutral protease, and DNase I at 37℃ for 30-45 min with gentle agitation. After digestion, the cell suspension was filtered through a 70-μm cell strainer, washed with cold PBS, and resuspended in staining buffer. When necessary, lamina propria mononuclear cells were further enriched by Percoll gradient centrifugation.

For surface staining, cells were stained with appropriate antibodies against surface antigens diluted in PBS on ice for 30 min. Cellular viability was assessed by staining with 7-aminoactinomycin D (7-AAD) (BioLegend, 420404) to exclude dead cells. For flow cytometric analysis, the cells were washed and resuspended in PBS. Flow cytometry was performed using a BD LSRFortessa flow cytometer (647800L6) controlled by FlowJo TM 10 software (BD, 647800L6). The single-cell suspensions were stained with the following antibodies: CD45-BV605 (BD, 563053), CD11b-PE (BioLegend, 101207), F4/80-BV786 (BioLegend, 123141), CD86-BV605 (BioLegend, 105125), CD86-PE/Dazzle-594 (BioLegend, 105041), CD206-FITC (BioLegend, 141703), CD206-APC (BioLegend, 141707).

**Caco-2 cells treatment by MEc30-NA**

For the MEc30-NA efficiency assay in Caco-2 cells, cells were grown to 80 to 90% confluence and then incubated for 12 h with serum-free medium (Control), supernatant of MEc30-WT, supernatant of MEc30-NA, and NA (50 μM) in the upper layer. Cells were collected after lysis by TRIzol reagent (Thermo Fisher, 15596018) lysate for qPCR.

**Wound-healing assay**

For the epithelial wound-healing assay, Caco-2 cells were seeded into 6-well plates and grown to full confluence. A linear scratch was generated in the monolayer using a sterile 200-μL pipette tip, followed by gentle washing with PBS to remove detached cells. The monolayers were then incubated for 24 h with serum-free medium (Control), supernatant of MEc30-WT, supernatant of MEc30-NA, or NA (50 μM). At the end of incubation, wound closure was imaged by phase-contrast microscopy, and the percentage of wound closure was quantified using ImageJ.

**Transwell permeability assay**

For the paracellular permeability assay, Caco-2 cells were seeded on collagen-precoated Transwell inserts (0.4-μm pore size) and cultured until formation of a polarized monolayer. Cells were then treated for 24 h with serum-free medium (Control), supernatant of MEc30-WT, supernatant of MEc30-NA, or NA (50 μM) added to the apical chamber. After treatment, FITC-dextran (4 kDa; Aladdin, F491425) was added to the apical compartment, and fluorescence intensity in the basolateral compartment was measured after 2 h as an indicator of epithelial permeability. Relative FITC-dextran flux was normalized to the Control group.

**cAMP Measurement**

Intracellular cAMP levels were quantified using a cAMP ELISA kit (Solarbio, SEKSM-0017) according to the manufacturer's instructions. Briefly, Caco-2 cells were seeded in 96-well plates and cultured to confluence. For Gi inhibition experiments, cells were pretreated with pertussis toxin (Sigma-Aldrich, P7208; 100 ng/mL) for 16 h prior to stimulation. Before the assay, cells were incubated with IBMX (Solarbio, 28822; 0.5 mM) for 15 min to inhibit phosphodiesterase activity. Subsequently, cells were treated with forskolin (Sigma-Aldrich, F6886; 10 μM) alone or in combination with nicotinic acid (NA, 50 μM) or bacterial supernatants from MEc30-WT or MEc30-NA for 15 min at 37 °C. Reactions were terminated by adding ice-cold 0.1 M HCl to lyse the cells and stabilize cAMP. Lysates were centrifuged, and the supernatants were neutralized before loading onto ELISA plates.

The cAMP content was determined by measuring absorbance at 450 nm and calculated from a standard curve generated using known cAMP concentrations. Data were normalized to the forskolin group and expressed as a percentage of the maximal response.

**Quantitative PCR**

For detection of gene expression in mouse tissues (including intestinal barrier-associated genes in the colon and inflammation-related genes in the liver), reverse transcription quantitative PCR (RT-qPCR) was performed. Extraluminal colon and liver tissues were rapidly excised, snap-frozen in liquid nitrogen, and stored at -80 °C until processing. Total RNA was isolated using TRIzol reagent (phenol–chloroform extraction), and 2 μg of RNA was reverse-transcribed into cDNA using a standard reverse transcription kit. RT-qPCR was carried out using the KAPA SYBR FAST qPCR Kit (10 μL of 2×Master Mix, 0.4 μL of each 10 μM forward and reverse primer, 1 μL of cDNA template, and nuclease-free water to a final volume of 20 μL). Amplification conditions were as follows: 95 °C for 3 min, followed by 40 cycles of 95 °C for 3 s, 60 °C for 20 s, and 72 °C for 20 s, after which a melting-curve analysis (65 °C for 5 s to 95 °C) was performed to verify amplicon specificity. Primer sequences are listed in Table S1. Gene expression was normalized to *Actb* (β-actin) and calculated using the $2^{-\Delta\Delta Ct}$

method, with values expressed as fold change relative to the control group. Standard curves were generated using serial dilutions of cDNA to confirm amplification efficiency.

For quantification of colonizing native *E. coli* strains in mouse fecal samples, genomic DNA was extracted from fecal pellets using a CTAB-based DNA extraction method. Briefly, each fecal pellet was resuspended in 600 μL CTAB lysis buffer and disrupted using a tissue lyser. The samples were then vortexed thoroughly and incubated at 65 °C for 1 h. After lysis, 700 μL chloroform-isoamyl alcohol (24:1) was added, followed by vortexing and centrifugation at 13,000 rpm for 15 min at room temperature. A total of 500 μL supernatant was transferred to a new 1.5 mL tube, mixed with 330 μL isopropanol by gentle inversion, and centrifuged at 12,000 × g for 2 min to precipitate genomic DNA. The DNA pellet was washed twice with 400 μL 70% ethanol, dried by vacuum centrifugation, and resuspended in 50 μL nuclease-free water. qPCR was then performed using the KAPA SYBR FAST qPCR Kit under the same cycling conditions as described above. For absolute quantification, standard curves were generated using serial dilutions of genomic DNA isolated from the corresponding native *E. coli* strain. Strain-specific primers targeting *E. coli*-GFP-specific genomic sequences were used to selectively detect the colonizing bacteria in fecal samples. Bacterial abundance was calculated based on Ct values and expressed relative to fecal input.

### ***In vivo*** **intestinal permeability assay**

Mice were fasted for 6 h then orally administered FITC-dextran 4 kDa (400 mg/kg body weight). After 90 min, 100 μL of blood were collected from the tip of the tail vein. The blood was kept in the dark and centrifuged at 3000×g for 10 min to collected plasma. Plasma aliquots (40 μL) were plated in 96-well plates and diluted to 200 μL with PBS. The fluorescence intensity in the plasma was measured at an excitation wavelength of 485 nm and an emission wavelength of 530 nm using a spectrofluorometer.

### **Immunohistochemistry**

Colon tissues were fixed in 4% paraformaldehyde in PBS, and the fixed sections were incubated in 3% $H_2O_2$ solution in PBS at room temperature for 10 min. Antigen retrieval was performed in sodium citrate buffer (0.01 M, pH 6.0) in a microwave oven at 1000 W for 3 min. Nonspecific antibody binding was blocked by incubation with 5% normal goat serum in PBS for 1 h at 25°C. Slides were stained overnight at 4°C with anti-CD86 antibody (CST, 19589S, 1:200 dilution) and anti-CD206 antibody

(CST, 24595S, 1:1000 dilution). The slides were subsequently washed and incubated with secondary antibodies for 30 min (HRP Goat Anti-Rabbit IgG (H+L), Abclonal, AS014). The sections were developed using a 3,3-Diaminobenzidine (DAB) substrate kit (ZSGB-BIO, ZLI-9017) and counterstained with hematoxylin. Images were captured using an Olympus BX600 microscope and a SPOT Flex camera. ImagePro Plus was used for further quantification of the DAB intensity in the image.

**Mass spectrometry detection for NA**

Quantification of different acylated NA was carried out using a LC-MS/MS system, consisting of an Acquity Ultra-Performance Liquid Chromatography (UPLC) system (Waters Corporation, Milford, USA) coupled to a Sciex 5500 triple quadrupole linear ion trap mass spectrometer (AB SCIEX, Framingham, MA, USA). Chromatographic separation was achieved using a Waters Xbridge HPLC C18 column (3.5 μm, 4.6×100 mm, Waters Corporation, Milford, USA) at 40°C, with a flow rate of 0.5 mL/min. The injection volume was 5 μL. Mobile phase A was 0.3% formic acid (FA) (Sigma-Aldrich, F0507) in water, and the mobile phase B was 0.3% FA in acetonitrile (Sigma-Aldrich, 34851): isopropanol (4:1). All analytes were detected in negative ion multiple reaction monitoring (MRM) mode. Chromatographic separation was performed with a linear gradient as follows: 0–2 min, 2% B; 2–4 min, 15% B; 4–5 min, 40% B; 5–7 min, 98% B; 7–9 min, 2% B. The detailed MRM parameters are listed in Table. Operational control of the LC-MS/MS system was performed using Analyst version 1.6.2, and quantitative analysis was carried out using MultiQuant software (version 3.0.1). The final data were processed, and concentrations were determined based on the standard curves and internal standard calibration.

| Metabolite | m/z | m/z | DP | CE |
|---|---|---|---|---|
| NA | 124 | 80 | 40 | 18 |

**Pharmacokinetics of NA/MEc30-NA**

MEc30-NA group mice were gavage with MEc30-NA engineered bacteria. After MEc30-NA engineered bacteria colonization was detected, mice in the NA group were gavaged with NA (500mg/kg) once, and then plasma was collected by inner canthus at 0,0.5,1,4,12 and 24 h time points. The levels of NA in mouse plasma were detected by LC-MS/MS.

**OGTTs and ITTs**

Mice were fasted for 6 h before the OGTTs and ITTs assays, as previously described[5]. The glucose levels were measured by tail vein blood sampling using a blood glucose test meter (Bayer HealthCare, Contour TS). After measurement of the fasting glucose levels, the mice were intraperitoneally injected with D-glucose (2 g/kg body weight; Sigma-Aldrich, G7021) for the GTTs or insulin (Sigma-Aldrich, 91077C; 1 IU/kg body weight) for the ITTs, and tail sampling was performed 15, 30, 60, and 90 min after the intraperitoneal injection for glucose level detection.

**Construction of *E. coli*-GFP, MEc30-Δ*clb*Δ*irp*, MEc30-mIL-10 and MEc30-NA**

GFP was inserted into the genome of EcN and other native *E. coli* at *exo* site, using the two-plasmid CRISPR-Cas9 system consisting of pEcCas (Addgene, 73227) and pEcgRNA derivatives. Taking EcN-GFP as an example, the dual-plasmid CRISPR-Cas9 system was used to integrate GFP into the exo locus of EcN. The guide RNA (gRNA) targeting the exo locus (5'-TTTATTGATATATTTACGTC-3') was designed using Benchling (https://benchling.com). The gRNA sequence was cloned into the pEcgRNA plasmid by PCR, generating pEcgRNA-exo. A repair template containing a 500 bp upstream homology arm, the GFP gene, and a 500 bp downstream homology arm was amplified by PCR and assembled into pEcgRNA-exo via overlap PCR to construct pTargetF-GFP. The pEcCas plasmid, which encodes Cas9 and the λ-Red recombination system, was electroporated into competent EcN cells to generate EcN-pEcCas. Subsequently, the pTargetF-GFP plasmid was electroporated into the resulting strain. To induce λ-Red recombination, cells were cultured in LB medium containing 10 mM arabinose at 37°C for 1 hour, followed by plating onto LB agar supplemented with kanamycin (50 μg/mL) and spectinomycin (50 μg/mL). Transformants were screened by colony PCR and DNA sequencing to confirm GFP insertion at the exo locus. To eliminate pTargetF-GFP, the positive clones were inoculated into LB medium containing 10 mM rhamnose and kanamycin (50 μg/mL) and incubated overnight at 220 rpm. The cultures were then diluted and plated onto LB agar containing kanamycin (50 μg/mL). Randomly selected colonies were transferred onto LB plates supplemented with kanamycin (50 μg/mL) and spectinomycin (50 μg/mL). Colonies that were sensitive to spectinomycin were considered cured of pTargetF-GFP. For the final plasmid curing step, the strain was inoculated into LB medium supplemented with 10 mM sucrose and cultured overnight at 30°C with shaking (220 rpm) to counter-select and eliminate the *sacB*-harboring pEcCas plasmid. No antibiotics were added during this step. The resulting cultures were then plated onto antibiotic-free LB agar, and single colonies were screened for plasmid loss by testing their sensitivity to kanamycin (50 μg/mL). Clones that failed to grow on LB agar containing kanamycin (50 μg/mL) but grew on antibiotic-free plates were selected as pEcCas-cured strains. The final recombinant strain was designated as EcN-GFP. The same editing workflow was subsequently applied to native *E. coli* isolates, including MEc30, following the EcN editing strategy, thereby generating GFP-tagged native strains for downstream experiments.

To generate the MEc30-Δ*clb*Δ*irp* strain, the *clbA-Q* and *irp1* loci were sequentially deleted from the MEc30 genome using the pEcCas/pEcgRNA two-plasmid CRISPR-Cas9 system. Briefly, gRNAs targeting the clb locus were designed and cloned into the pEcgRNA plasmid. Repair templates containing the upstream and downstream homology arms flanking the target region were amplified by PCR and assembled into the corresponding targeting plasmid. The pEcCas plasmid was first electroporated into competent MEc30 cells, followed by electroporation of the corresponding targeting plasmid. After λ-Red induction, transformants were selected on LB agar containing kanamycin and spectinomycin. The *clb* locus was deleted as a first step, as previous studies have shown that removal of the entire pks island (*clbA*–*clbQ*) is sufficient to abolish colibactin production without impairing the growth or colonization capacity of EcN[6]. Plasmid curing was performed as described above. Subsequently, the *irp* locus was deleted in the MEc30-Δ*clb* background using the same workflow, including gRNA construction, repair template assembly, electroporation, λ-Red induction, and antibiotic selection. This design was based on previous evidence showing that *irp1* deletion can be used to generate a yersiniabactin-deficient strain. The resulting double-knockout strain was designated MEc30-Δ*clb*Δ*irp*. For genotyping, the Δ*clb* deletion was verified using locus-specific validation primers. Positive Δ*clb* colonies yielded an approximately 1.4 kb PCR product, whereas the wild-type allele did not generate a detectable band under the same amplification conditions because the expected fragment was too large. The Δ*irp* deletion was similarly verified using locus-specific validation primers, with positive Δ*irp* colonies yielding an approximately 1.2 kb PCR product, while the wild-type allele showed no detectable band for the same reason. All subsequent engineered strains were constructed on the basis of MEc30-ΔclbΔirp. Thus, strains such as MEc30-mIL-10 and MEc30-NA were formally derived from MEc30-ΔclbΔirp-mIL-10 and MEc30-ΔclbΔirp-NA, respectively, but are abbreviated throughout the manuscript as MEc30-mIL-10 and MEc30-NA for simplicity.

The construction of MEc30-mIL-10 and MEc30-NA was based on integration into the *attB* site using the pTargetF plasmid pTarget-Dpp4, as described in our previously published study[7]. Building upon this, we designed primers to generate pTarget-mIL-10 and pTarget-NA. Details are provided in Supplementary Table 1. Specifically, MEc30-mIL-10 was constructed by integrating a murine IL-10 expression cassette into the *attB* site of the MEc30-GFP genome using a two-plasmid CRISPR–Cas9 system. The cassette contained the murine IL-10 coding sequence fused to the bacterial secretion signal peptide PelB and driven by the constitutive HCE promoter, a high-level expression promoter derived from the upstream regulatory region of the D-amino acid aminotransferase gene of *Geobacillus toebii*, which enables inducer-free and continuous transgene expression in E. coli. Following CRISPR-mediated double-strand cleavage, the cassette was inserted via λ-Red homologous recombination[8]. Correct integration of the mIL-10 cassette at the *attB* site was confirmed by colony PCR using primers flanking the recombination junctions, which produced an extra 700-bp fragment in MEc30-mIL-10.

To enable endogenous NA biosynthesis and construct MEc30-NA, the *pyrZ* gene was integrated into the genome of the native colonizing strain MEc30. The integration site was the chromosomal *attB* locus, which provides a neutral genomic region for stable gene insertion without affecting bacterial growth or fitness. The *pyrZ* gene encodes a bifunctional NadC homolog previously characterized as a key enzyme catalyzing nicotinic acid formation during pyridomycin biosynthesis. The *pyrZ* sequence used in this study was derived from *Streptomyces pyridomyceticus* and was commercially synthesized after codon optimization for *E. coli* expression[9]. A constitutive promoter was used to drive stable *pyrZ* expression. Correct integration of the *pyrZ* cassette at the *attB* site was confirmed by colony PCR using primers flanking the recombination junctions, which produced an extra 1,000-bp fragment in MEc30-NA.

All PCR products were confirmed by sequencing to validate the correct gene insertions.

**QUANTIFICATION AND STATISTICAL ANALYSIS**

GraphPad Prism (version 9.0) and SPSS (version 27.0) were used for statistical analysis. Experimental data are presented as the mean ± SEM. The sample size was determined based on previous experience, sample availability, and prior studies. No data were excluded from the analysis. The normality of the data distribution was assessed using the Shapiro-Wilk test. For statistical comparisons, Student's t-test was used for two-group comparisons, while one-way ANOVA was applied for multiple-group comparisons, followed by Tukey's post hoc test (for equal standard deviations) or Dunnett's T3 test (for unequal standard deviations). For non-normally distributed data, the Mann-Whitney U test was used for two-group comparisons, and the Kruskal-Walli's test was applied for multiple-group comparisons. A *p*-

value $<0.05$ was considered statistically significant.
